# Prolactin-mediates a lactation-induced suppression of arcuate kisspeptin neuronal activity necessary for lactational infertility in mice

**DOI:** 10.1101/2024.01.26.577359

**Authors:** Eleni C.R. Hackwell, Sharon R. Ladyman, Jenny Clarkson, H. James McQuillan, Ulrich Boehm, Allan E. Herbison, Rosemary S.E. Brown, David R. Grattan

## Abstract

The specific role that prolactin plays in lactational infertility, as distinct from other suckling or metabolic cues, remains unresolved. Here, deletion of the prolactin receptor (Prlr) from forebrain neurons or arcuate kisspeptin neurons resulted in failure to maintain normal lactation-induced suppression of estrous cycles. Kisspeptin immunoreactivity and pulsatile LH secretion were increased in these mice, even in the presence of ongoing suckling stimulation and lactation. GCaMP fibre photometry of arcuate kisspeptin neurons revealed that the normal episodic activity of these neurons is rapidly suppressed in pregnancy and this was maintained throughout early lactation. Deletion of Prlr from arcuate kisspeptin neurons resulted in early reactivation of episodic activity of kisspeptin neurons prior to a premature return of reproductive cycles in early lactation. These observations show dynamic variation in arcuate kisspeptin neuronal activity associated with the hormonal changes of pregnancy and lactation, and provide direct evidence that prolactin action on arcuate kisspeptin neurons is necessary for suppressing fertility during lactation in mice.

## Introduction

In mammals, lactation is accompanied by a period of infertility. This adaptive change establishes appropriate birth spacing to enable maternal metabolic resources to be directed towards caring for the new-born offspring, rather than supporting another pregnancy (1). Lactational infertility is characterized by a lactation-induced suppression of pulsatile luteinizing hormone (LH) secretion, and the temporary loss of the reproductive cycle (in rodents this is exhibited as an extended period of diestrus or anestrus) (2–5). Lactation is also characterised by chronically elevated levels of the anterior pituitary hormone prolactin, which is essential for milk production and promotes adaptive changes in maternal physiology and behaviour (1,2,4,5). Despite hyperprolactinaemia being a well-recognized cause of infertility, the specific role that prolactin plays in lactational infertility, as distinct from other suckling- or metabolic-related cues, is currently unclear (4,6).

Recent *in vivo* studies have confirmed that kisspeptin neurons in the arcuate nucleus of the hypothalamus are responsible for the periodic release of gonadotrophin-releasing hormone (GnRH) and subsequent pulsatile LH secretion that drives reproductive function (7–11). Studies using GCaMP fibre photometry in conscious mice have demonstrated that the arcuate kisspeptin neuronal population exhibits episodes of increased intracellular calcium levels coincident with, and immediately preceding, each pulse of LH secretion in intact and gonadectomised male and female mice (7–9). Miniscope investigations showed that individual kisspeptin neurons within the arcuate population act in a coordinated, synchronised, and episodic manner (10,11). Loss of pulsatile LH secretion during lactation and consequent lactational infertility may be caused by the loss of kisspeptin-mediated stimulation of GnRH secretion (12–18). Kisspeptin expression is markedly suppressed in lactation (12,16) and even when exogenously stimulated, kisspeptin neurons are unable to activate GnRH neurons during lactation, likely due to a lack of kisspeptin synthesis (13).

It is well established that hyperprolactinemia causes infertility, and thus, the elevated prolactin present in lactation seems a likely candidate to be involved in suppressing fertility during lactation. Prolactin administration acutely suppresses LH secretion (19), and chronic exposure to elevated prolactin reduces *Kiss1* mRNA expression in the arcuate nucleus (17,20,21). In lactating mice, suppressing endogenous prolactin secretion shortens the period of infertility (22), suggesting that prolactin is important for maintaining the suppression of pulsatile LH secretion during lactation. Such a role for prolactin is controversial, however (4,6,23–27), with studies in a number of species suggesting that the neural stimulation of suckling may be more important than prolactin in maintaining lactational infertility (28,29). However, it has previously been difficult to disentangle the specific role of prolactin, as suckling, prolactin, and milk production are so tightly linked that manipulating one ultimately impacts the others, making it difficult to determine the contribution of any one element. Here, using a conditional deletion strategy, we have blocked prolactin action in the brain leaving suckling, lactation, and maternal behaviour intact. Using GCaMP fibre photometry techniques, we have also documented arcuate kisspeptin neuron activity across pregnancy and lactation transitions in the same mice and established that prolactin directly acts on these neurons to suppress fertility in lactation.

## Methods

### Animals

All experiments were performed using adult female mice on a C57BL/6J background (8-20 weeks of age unless otherwise stated). Mice were housed under controlled temperature (22 °C ± 2°C) and lighting (12-hour light/12-hour dark schedule, with lights on at 0600 hours) with *ad libitum* access to food and water (Teklad Global 18% Protein Rodent Diet 2918; Envigo, Huntingdon, United Kingdom). Daily body weight was recorded and daily vaginal cytology was used to monitor the estrous cycle stage. All experiments were carried out with approval from the University of Otago Animal Welfare and Ethics Committee.

To establish pregnancies, mice were mated with male wild-type C57BL/6J mice (presence of sperm plug = day 1 pregnancy). Male mice were then removed once a sperm plug was seen, and no male mice were present at parturition. The first day a litter was seen was counted as day 1 of lactation and maternal mice were left undisturbed till day 3 of lactation, when vaginal monitoring would resume and litter size was normalised to 6 pups per animal, unless otherwise stated. The intention of the normalisation was to reduce variability in the degree of suckling stimulus each dam would receive.

Towards the end of each recording period, bilateral ovariectomy (OVX) was performed under isoflurane anaesthesia with pre- and post-operative Carprofen (5mg/kg, s.c.) administered for pain relief. This enabled us to examine post-OVX changes in activity of kisspeptin neurons and expression of kisspeptin in the arcuate nucleus.

### Pulsatile hormone measurement

To monitor pulsatile secretion of LH, serial tail tip blood sampling and measurement of LH by ELISA was undertaken as reported previously (19,30,31). As novel exposure and restraint stress has been shown to suppress pulsatile LH secretion (32), all mice were habituated to the tail tip blood sampling procedure by picking up the mouse in a gentle restraint device (soft cardboard tube) or hand and lightly massaging the tail for approximately 5 minutes per day for at least 3 weeks prior to experimentation (33). Sequential whole blood samples (4μl) were collected in 6 minute intervals for 3 hours between 0900 and 1200 hours, unless otherwise stated. Samples were immediately diluted in 48ul 0.01M PBS/0.05% Tween 20, and frozen on dry ice before being stored at -20°C for subsequent LH measurement.

### RNAscope

Mice were transcardially perfused with a micro-perfusion pump with 2% paraformaldehyde (PFA). Brain sections (14μm-thick) were prepared, thaw mounted onto superfrost-plus microscope slides and then stored at -80°C. RNAscope in-situ hybridization was performed using the RNAscope 2.5 High definition Duplex Detection kit – chromogenic (Advanced Cell Diagnostics, Hayward, CA) largely in accordance with manufacturer’s instruction. The channel 1 Prlr probe was custom designed to pick up only the long form of the prolactin receptor. It was designed to transcript NM_011169.5 with a target sequence spanning nucleotides 1107-2147 (Ref: 588621; Advanced Cell Diagnostics, Hayward, CA). The channel 2 Kiss1 probe was custom designed to transcript NM_178260.3 with a target sequence spanning nucleotides 5 to 485 (Ref: 500141-C2; Advanced Cell Diagnostics, Hayward, CA). Sections were thawed at 55°C, postfixed for 3 minutes in 2% PFA, washed in 0.01M PBS for 5 minutes, and endogenous peroxidases were blocked with a hydrogen peroxidase solution for 10 minutes. Tissue was washed in distilled water (3x 2 minutes), then briefly immersed in 100% ethanol, air dried for 5 minutes, and a hydrophobic barrier was applied. Tissue was permeabilized with RNAscope protease plus for 30 minutes at 40°C. Sections were washed (2 x 2 minutes) and were hybridized with the Prlr and Kiss1 probes (1:300 dilution, Prlr:Kiss1) or negative control probe (Cat#320751; Advanced Cell Diagnostics, Hayward, CA) at 40°C for 2 hours. Amplification (Amp 1-6) was performed in accordance with the manufacturer’s instructions. Sections were then hybridized with a Fast-RED (1:60, Fast-RED B:Fast-RED A) for 10 minutes at room temperature, before undergoing further amplification steps (Amp 7-10) in accordance to manufacturer’s instructions. The final positive hybridization was detected by incubation with the secondary detection reagents (1:50, Fast-GREEN B:Fast-GREEN) for 10 minutes at room temperature. Sections were washed, counterstained with haematoxylin (25% Gills), dried at 60°C for 20 minutes, and cover-slipped with VectaMount (Vector laboratories, H-5000) before imaging using an Olympus BX51 light microscope and Olympus UPlanSApo 10/20x lenses.

Quantification of the proportion of kisspeptin neurons co-expressing *Prlr* mRNA was undertaken in FIJI software (National Institute of Health, Bethesda, Maryland, USA) following image acquisition. Positive hybridization for *Kiss1* and *Prlr* was determined based on scoring guidelines included in the manufacturer’s protocol (Advanced Cell Diagnostics). The total number of *Kiss1*-expressing cells and the total number of these that showed *Prlr* mRNA expression were counted.

### Effect of neuron-specific deletion of the prolactin receptor gene on the maintenance of lactational infertility

To investigate whether prolactin action in the brain is required for lactational infertility, neuron-specific *Prlr* knockout mice (*Prlr*^lox/lox^/*Camk2a*^Cre^) and their respective Cre-negative controls (*Prlr*^lox/lox^) were generated (as previously described in (34)). We have previously shown that while *Prlr*^lox/lox^/*Camk2a*^Cre^ mice do not have a complete Prlr deletion in the forebrain. There are areas of extensive deletion (as measured by reduced prolactin-induced pSTAT5), such as the arcuate nucleus and ventromedial nucleus of the hypothalamus, and there are also areas where Prlr is reduced by about 50% such as the medial pre-optic area (34,35). In our experience with these animals, this is sufficient to retain normal maternal behaviour in most animals (our unpublished data), and this was the case for the animals used in the present study. RNAscope in-situ hybridization was done to confirm the degree of knockout in intact, diestrous, nulligravid, non-lactating (henceforth termed NL) mice and 14 day post-OVX mice (*Prlr*^lox/lox^ intact n = 6, *Prlr*^lox/lox^ OVX n = 5, *Prlr*^lox/lox^/*Camk2a*^Cre^ intact n = 5, *Prlr*^lox/lox^/*Camk2a*^Cre^ OVX n = 5; all aged 8-16 weeks). Both intact and OVX mice were included as kisspeptin cell bodies are only visible in the rostral periventricular region of the third ventricle (RP3V) of intact mice and in the arcuate nucleus of OVX mice, due to estradiol regulation (36). *Prlr*^lox/lox^/*Camk2a*^Cre^ mice showed a significant decrease in the percentage of *Kiss1*-expressing cells co-expressing *Prlr* compared to controls in both the RP3V (p = <0.0001) and arcuate nucleus (p = 0.0009) (unpaired two-tailed t tests, Supplementary Figure 1A-D).

*Prlr*^lox/lox^/*Camk2a*^Cre^ mice are hyperprolactinaemic due to impaired negative feedback of prolactin on hypothalamic dopamine neurons (34) and therefore show disrupted estrous cycles (showing recurrent pseudopregnancy-like cycles with long periods of diestrus of approximately 14 days between estrous stages). However, these mice are able to become pregnant and have normal pregnancies. All *Prlr*^lox/lox^/*Camk2a*^Cre^ mice were given a 250μl subcutaneous injection of bromocriptine (5mg/kg, 5% ethanol/saline; Tocris Bioscience Cat#0427) prior to being mated. This treatment was designed to reinstate an estrous cycle in *Prlr*^lox/lox^/*Camk2a*^Cre^ mice. Bromocriptine is an agonist for the type 2 dopamine receptor and inhibits prolactin secretion from the pituitary gland (37), thereby terminating the pseudopregnancy-like state and bringing the mice into proestrus the following day. Following treatment, all mice were then housed with a stud male.

For *Prlr*^lox/lox^/*Camk2a*^Cre^ mice, estrous cycles were monitored from day 3 of lactation until the first day of diestrus following a day of estrus (proestrus and estrus had to be observed prior to transcardial perfusion on the first day of diestrus). Brains were collected following transcardial perfusion for assessment of kisspeptin immunoreactivity. For every lactating *Prlr*^lox/lox^/*Camk2a*^Cre^ mouse (n = 8), the brain of a *Prlr^l^*^ox/lox^ control mouse (n = 8) of the equivalent day (±1) of lactation was also collected. A group of NL mice of both genotypes (n = 5-6) was also perfused for immunohistochemistry on diestrus.

To evaluate pulsatile LH secretion in early lactation (prior to the return of estrous cycles) and to determine whether progesterone played any role in regulating pulsatile LH secretion in lactation, additional groups of lactating *Prlr*^lox/lox^/*Camk2a*^Cre^ and *Prlr^l^*^ox/lox^ control mice were generated. These mice were treated with either the progesterone receptor antagonist, mifepristone (4mg/kg in sesame oil, s.c.; AK Scientific Inc Cat#J10622), or vehicle (n = 7-8 per group) on the morning of day 4 of lactation and day 5 of lactation. This dose was selected as it was found to be sufficient to cause termination of pregnancy in wild-type C57BL/6J mice (p = 0.0072, chi-squared test, Supplementary Figure 2A; pilot study, n = 6 both groups). Neither vehicle nor mifepristone treatment had an effect on litter weight gain (interaction of time x genotype & treatment p = 0.5322, two-way repeated measures ANOVA, Supplementary Figure 2B). Mice underwent blood sampling for LH for 3 hours on day 5 of lactation, 30 minutes after treatment.

### Measurement of LH concentrations

An established sandwich ELISA method was used to determine LH concentration in diluted whole blood samples collected from mice (30,31). Briefly, a 96-well high plate was incubated with bovine monoclonal antibody (LHβ518b7, 1:1000 in 1xPBS; Dr. L. Sibley, UC Davis, CA, USA) for 16 h at 4°C. Following incubation of standards, controls and experimental samples for 2 hours, plates were incubated in rabbit polyclonal LH antibody (AFP240580Rb; 1:10,000; National Hormone and Pituitary Program, NIH) for 90 minutes, followed by incubation with polyclonal goat anti-rabbit IgG/HRP antibody (1:1000; DAKO Cytomation) for 90 minutes. Finally, plates were incubated in OPD (o-phenylenediamine capsules; Sigma-Aldrich Cat#P7288) for 30 minutes. A standard curve for the detection of LH concentration was generated using serial dilutions of mouse LH-reference preparation peptide (National Hormone and Pituitary Program, NIH). LH levels were read using a standard absorbance plate reader (SpectraMax ABS Plus; Molecular Devices) at 490nm and 630nm wavelengths. The assay had a sensitivity of 0.04ng/ml to 4ng/ml, with an intra-assay coefficient of variation of 4.40% and an inter-assay coefficient of variation of 8.29%.

PULSAR Otago was used to define LH pulses (38). Parameters used; Smoothing 0.7, Peak split 2.5, Level of detection 0.04, Amplitude distance 3, Assay variability 0, 2.5, 3.3, G(1)=3.5, G(2)=2.6, G(3)=1.9, G(4)=1.5, G(6)=1.2. Mean LH levels were calculated by averaging all LH levels collected during the experiment. Individual LH profiles from mice are shown in Supplementary Figure 3.

### Assessment of kisspeptin expression

#### Perfusion and fixation of tissue

Mice were anaesthetised with sodium pentobarbital (15mg/mL, i.p.) and transcardially perfused with 4% PFA. Brains were removed, postfixed in the same solution, and cryoprotected overnight in 30% sucrose before being frozen at -80°C. Two sets of 30μm thick coronal brain sections were cut using a sliding microtome, from Bregma 1.10mm to - 2.80mm. Brain sections were kept in cryoprotectant solution (pH = 7.6) at -20°C until immunohistochemistry was performed.

#### Immunohistochemistry

Immunohistochemistry for kisspeptin in the RP3V and arcuate nucleus was performed as previously described (39). Briefly, sections were incubated in polyclonal rabbit anti-kisspeptin primary antibody (AC 566, 1:10,000; gift from A. Caraty, Institut National de la Recherche Agronomique, Paris, France) for 48 hours at 4°C. Sections were then incubated with biotinylated goat anti-rabbit IgG (1:200, Vector biolabs, Peterborough, GK) for 90 min at room temperature, followed by incubation in an avidin-biotin complex (Elite vectastain ABC kit, Vector laboratories). The bound antibody-peroxidase complex was visualised using a nickel-enhanced diaminobenzidine (DAB) reaction, to form a black cytoplasmic precipitate.

Brain sections were imaged as described in RNAscope section above. Quantification of kisspeptin neurons in the RP3V, was undertaken by manually counting all labelled neurons present in all three subdivisions: the anteroventral periventricular nucleus (AVPV); rostral preoptic periventricular nucleus (rPVpo); and caudal preoptic periventricular nucleus (cPVpo). Two sections per subdivision per mouse were averaged for each animal. As kisspeptin cell bodies in the arcuate nucleus were not easily observed, as previously reported (40), kisspeptin fibre immunoreactivity was imaged using a Gryphax NAOS colour camera (Jenoptik) and evaluated using FIJI software and the voxel counter function (National Institutes of Health). Kisspeptin fibre density was measured in the arcuate nucleus across the three subdivisions; rostral arcuate (rARC), middle arcuate (mARC), and caudal arcuate (cARC). Two sections of each subdivision per animal were counted and then averaged across each animal to get total number and reported as total amount of voxels per ROI (voxel fraction).

### Characterization of arcuate kisspeptin neuronal activity using GCaMP fibre photometry

#### Stereotaxic surgery and AAV injections

Adult *Kiss1*^Cre^ or *Prlr*^lox/lox^/*Kiss1*^Cre^ mice (2-3 months old) were anaesthetised with 2% Isoflurane, given local Lidocaine (4mg/kg, s.c.) and Carprofen (5mg/kg, s.c.) and placed in a stereotaxic apparatus. A custom-made unilateral Hamilton syringe apparatus holding one Hamilton syringe was used to perform unilateral injections into the arcuate nucleus. The needles were lowered into place (-0.14mm A/P, +0.04mm M/L, -0.56mm DV) over 2 minutes and left in situ for 3 minutes before injection was made. 1μl AAV9-CAG-FLEX-GCaMP6s-WPRE-SV40 (1.3x10^-13^ GC/ml, University of Pennsylvania Vector Core, Philadelphia, PA, USA) was injected into the arcuate nucleus at a rate of ∼100nl/min with the needles left in situ for 3 minutes prior to being withdrawn over a period of 6 minutes. This was followed by implantation of a unilateral indwelling optical fibre (400 μm diameter, 6.5 mm long, 0.48 numerical aperture (NA), Doric Lenses, Canada, product code: MFC_400/430-0.48_6.5mm_SM3*_FLT) at the same coordinates. Carprofen (5mg/kg body weight, s.c.) was administered for post-operative pain relief. After surgery, mice received daily handling and habituation to the fibre photometry recording conditions over 4-6 weeks before experimentation began.

#### GCaMP fibre photometry

Photometry was performed as reported previously (9). Fluorescence signals were acquired using a custom-built fibre photometry system made primarily from Doric components. Violet (405nm) and blue (490nm) fibre-coupled LEDs were sinusoidally modulated at 531 and 211 Hz, respectively, and focused into a 400μm, 0.48 numerical aperture fibre optic patch cord connected to the mouse. Emitted fluorescence was collected by the same fibre and focused onto a femtowatt photoreceiver (2151, Newport). The two GCaMP6s emission signals were collected at 10 Hz in a scheduled 5s on/15s off mode by demodulating the 405nm (non-calcium dependent) and 490nm (calcium dependent) signals. The power output at the tip of the fibre was set at 50μW. Fluorescent signals were acquired using a custom software acquisition system (Tussock Innovation, Dunedin, New Zealand) and analysed using custom templates created by Dr Joon Kim (University of Otago, Dunedin, New Zealand) based on mathematics and calculations similar to those previously described (41,42). Briefly, the fluorescent signal obtained after stimulation with 405nm light was used to correct for movement artefacts as follows: first, the 405nm signal was filtered using a savitzky-golay filter and fitted to the 490nm signal using least linear square regression. The fitted 405nm signal was then subtracted and divided from the 490nm signal to obtain the movement and bleaching corrected signal. The output of these templates is 490-adjusted405/adjusted405, which was multiplied to get the final ΔF/F as a percentage increase. All photometry data is reported as ΔF/F(%).

All recordings were obtained from freely behaving mice between 0800 hours and 1200 hours (apart from 24 hours post weaning recording (0900 hours to 1700 hours), and day 18/19 pregnancy recording (1800 hours to 0800 hours the following day)). Synchronized events (SEs) were defined as when ΔF/F exceeds 3 standard deviations (SD) above the trace mean. Manual SE shape analysis was performed in addition to standard deviation method for certain datasets where necessary. SEs were counted manually to determine frequency of SEs per 60 minutes. The between-animal variability in total signal means that changes in SE amplitude can only be reported as relative changes within an animal. Relative SE amplitude was calculated by using normalised ΔF/F data and then subtracting the peak of an SE from the nearest nadir to the rise of the SE and averaging that for the number of SEs in a recording. To obtain normalised ΔF/F, three pre-pregnancy datasets from each mouse were used to find the average maximum ΔF/F for that mouse. All datasets were then divided by that normalisation value to get normalised ΔF/F for each trace for each individual mouse. In these long-term longitudinal studies, some data points are missing due to issues with recordings (e.g. dirty fibre optic cannula or system battery issues), meaning that occasionally fewer mice are used for analysis at various time points. During experimentation, no mice lost cannulas, but a gradual reduction in amplitude of signal was observed over time.

Following experimentation, brains were collected in a similar way to that described above. Postmortem analysis of cannula placement and GCaMP6 transfection in each animal was not done, because with experience, the quality and characteristics of the SE recordings (including corresponding pulsatile LH secretion) in each animal was indicative of successful cannula placement and transfection of the GCaMP. In a preliminary trial the green fluorescent protein (GFP) attached to the GCAMP was restricted to the caudal arcuate nucleus (n = 3 mice, preliminary trial), with little evidence of spread into the rostral or middle arcuate. GFP was seen exclusively ipsilaterally, with no GFP observed on the contralateral side. Cannula placement in these trial animals was in the caudal arcuate.

#### Monitoring the activity of arcuate kisspeptin neurons across different reproductive stages in the same mice

Adult *Kiss1*^Cre^ mice were 8-10 weeks of age at the beginning of experiments, and up to 12 months old by the time of final recording (n = 8 during pregnancy; n = 6 during lactation, 2 mice were euthanised due to dystocia). Monitoring of vaginal cytology and weight was continuous from 1 week pre-surgery till day 19 of pregnancy and resumed on day 3 of lactation until the end of experiment. All handling stopped on day 19 of pregnancy to avoid potential compromise of parturition and onset of maternal behaviour.

To investigate the activity of the arcuate kisspeptin population across different reproductive states in the same animal, the following recording protocol was followed for all *Kiss1*^Cre^ mice, unless otherwise stated; diestrus NL, day 4 of pregnancy, day 14 of pregnancy, day 18/19 of pregnancy (overnight), day 7 of lactation, day 14 of lactation, day 18 of lactation, 24 hours after weaning, first diestrus after estrous cycles begin following weaning, and 10 days post-OVX. In addition, blood sample collection for paired LH measurement was done in diestrus NL state, on day 14 of pregnancy (in 4 mice, maximum of 6 samples were collected around small “peaks” in baseline), day 7 of lactation, and day 14 of lactation. During sampling, blood samples were taken at the usual 6 minute intervals previously described, however if a SE was observed and the next scheduled sampling was more than 3 minutes away, an extra blood sample was taken 2 minutes after the SE. Following this additional sampling, the next scheduled blood sample would be taken at its original time and then every 6 minutes after, till another SE was seen. Blood sampling was not carried out at additional time points as blood sampling was undertaken at least a week apart, and stress from repeated sampling was attempted to be kept at a minimum e.g., not blood sampling around the time of birth. Fibre photometry recordings were usually between 2-4 hours in length. The only longer recordings were undertaken on day 18/19 of pregnancy (14 hours) and 24 hours after weaning (8 hours). These particular recording sessions were extended to determine whether there were any longer-term changes occurring in the activity of the arcuate kisspeptin population in the lead up to parturition or following weaning of pups (states closely followed by postpartum estrus and resumption of normal estrous cycles, respectively).

#### Effect of arcuate kisspeptin neuron-specific deletion of the prolactin receptor gene on the maintenance of lactational infertility and the activity of kisspeptin neurons during lactation

Kisspeptin-specific prolactin-receptor knockout mice (*Prlr*^lox/lox^/*Kiss1*^Cre^) and their respective Cre-negative controls (*Prlr*^lox/lox^) were generated as previously described (19). *Prlr*^lox/lox^/*Kiss1*^Cre^ (n = 27) and *Prlr*^lox/lox^ control (n = 30) dams underwent estrous cycle monitoring from day 3 of lactation onwards to determine whether mice showed an early resumption of estrous cycles. RNAscope in-situ hybridization (as described above) was done to confirm knockdown in a subgroup of these mice, with *Prlr*^lox/lox^/*Kiss1*^Cre^ mice showing a significant decrease in the percentage of *Kiss1*-expressing cells co-expressing *Prlr* compared to controls in the arcuate nucleus (p = <0.0001, unpaired two-tailed t test, Supplementary Figure 1E-G; *Prlr*^lox/lox^ n = 6, *Prlr*^lox/lox^/*Kiss1*^Cre^ n = 8).

To determine whether the deletion of the prolactin receptor from arcuate kisspeptin neurons led to early reactivation of these neurons during lactation, adult *Prlr*^lox/lox^/*Kiss1*^Cre^ mice (n = 5) and an additional *Kiss1*^Cre^ control mouse (n = 1) (8 weeks old at the start of the experiment, and up to 14 months at final recording timepoint) were set up for fibre photometry, as described above. Monitoring of vaginal cytology and weight was continuous from 1 week pre-surgery till day 18 of pregnancy and resumed on day 3 of lactation. Recordings were undertaken in a similar timeline to that described above, however no pregnancy recordings were done, and in early lactation recordings were performed every 2 days from day 3 to day 9 of lactation, before following the same protocol as described. No blood samples were taken over the lactation period in these mice. As described previously, recordings were kept between 2-4 hours, apart from the 24 hours after weaning recording (8 hours).

### Statistical analysis

Data are presented as mean ± SEM and all statistical analysis was performed with PRISM software 10 (GraphPad Software, San Diego, CA, USA) with a p value of < 0.05 considered as statistically significant. Individual symbols in graphs represent individual mice. Differences in kisspeptin cell number or fibre density was assessed using two-way ANOVAs with Tukey’s multiple comparisons tests or t tests, with both analyses using combined averages of each animal (averaged number of cells or fibre density across the three subdivisions of each nucleus to get total number reported). Resumption of estrous cycles was analysed using Log-rank (Mantel-Cox) test chi-squared test. LH pulse frequency data and mean LH data was analysed using two-way ANOVAs with Tukey’s multiple comparisons tests. SE frequency and amplitude throughout reproductive cycles was analysed using mixed effect analysis (fixed type III) with Tukey’s multiple comparisons tests where appropriate and day 18/19 of pregnancy data was analysed using t tests. Correlation between SE occurrence and LH pulses was assessed using chi-squared test. All fibre photometry data used for quantitative analysis and comparison were from the first pregnancy and lactation. A full list of probability values, inferential statistics, and degrees of freedom for all data can be found in Supplementary Table 1.

## Results

### Prolactin action on forebrain neurons is necessary to maintain lactational infertility

Lactation has previously been associated with a marked decrease in *Kiss1* mRNA levels in both RP3V and arcuate nucleus populations (13). To determine whether prolactin was involved in the maintenance of lactational anestrus, the *Prlr* gene was knocked out of *Camk2a* expressing neurons (most forebrain neurons) of female mice. Control *Prlr*^lox/lox^ mice showed a marked reduction in kisspeptin cell body immunoreactivity in the RP3V of lactating compared to NL mice (p = 0.0100, *Post hoc* Tukey’s multiple comparisons test, Figure 1A, C). In contrast, in *Prlr*^lox/lox^/*Camk2a*^Cre^ mice the lactation-induced suppression of kisspeptin cell bodies in the RP3V was absent (p = 0.6409, *Post hoc* Tukey’s multiple comparisons test; interaction between reproductive state and genotype p = 0.0034, two-way ANOVA, Figure 1A, C). Fibre density in the arcuate nucleus was significantly increased in lactating *Prlr*^lox/lox^/*Camk2a*^Cre^ mice compared to lactating *Prlr*^lox/lox^ controls (p = 0.0020, unpaired two tailed t test, measured as percentage voxels within the region of interest, Figure 1B, D).

**Figure 1.**
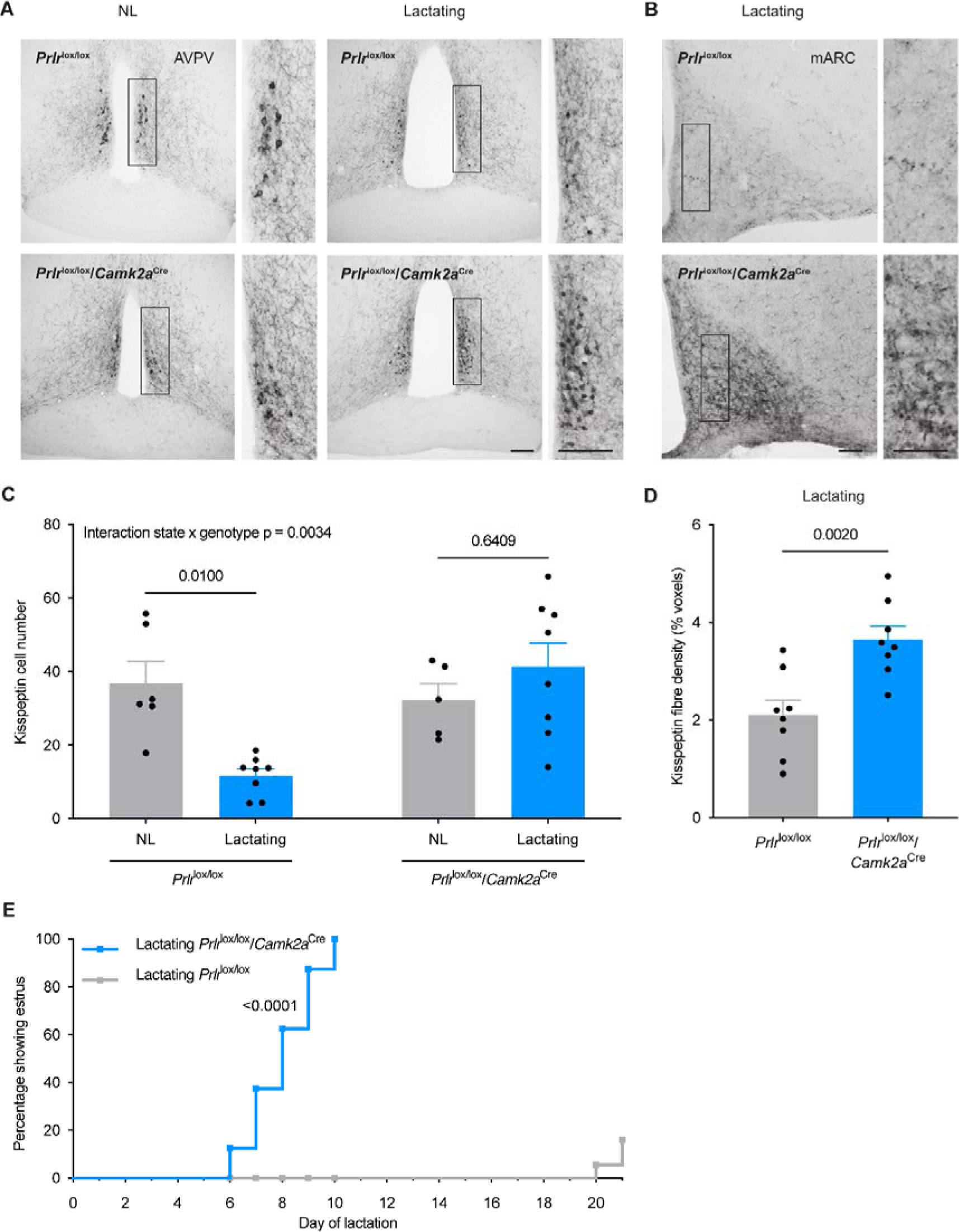
*Prlr*^lox/lox^/*Camk2a*^Cre^ mice do not undergo the normal period of lactational infertility and the lactation-induced suppression of kisspeptin immunoreactivity is absent. (A) Kisspeptin immunoreactivity shown in representative photomicrographs from diestrus, nulligravid, non-lactating (NL; left) and lactating (right) *Prlr*^lox/lox^ control and *Prlr*^lox/lox^/*Camk2a*^Cre^ mice (from anteroventral periventricular nucleus (AVPV) region of RP3V). (B) Representative photomicrographs showing mid arcuate nucleus (mARC) of lactating *Prlr*^lox/lox^ (top) and lactating *Prlr*^lox/lox^/*Camk2a*^Cre^ mice (bottom). (C) Total kisspeptin cell number for the RP3V (NL *Prlr*^lox/lox^ (n = 6) versus lactating *Prlr*^lox/lox^ control (n = 8) p = 0.0100, NL *Prlr*^lox/lox^/*Camk2a*^Cre^ (n = 5) versus lactating *Prlr*^lox/lox^/*Camk2a*^Cre^ (n = 8) p = 0.6409). Two-way ANOVA followed by Tukey’s multiple comparisons test. (D) Quantification of kisspeptin fibre density in the arcuate nucleus (Fiji software, measured in percentage voxels per region of interest), showing total kisspeptin fibre density in the arcuate nucleus (lactating *Prlr*^lox/lox^ control n = 8, lactating *Prlr*^lox/lox^/*Camk2a*^Cre^ n = 7, p = 0.0020, unpaired two-tailed t test). (E) Lactating *Prlr*^lox/lox^/*Camk2a*^Cre^ mice (blue, n = 8) resume estrous cycles significantly earlier (100% within 6-10 days of lactation) than lactating *Prlr*^lox/lox^ controls (grey, n = 10) (p = <0.0001, Log-rank (Mantel-Cox) test). Scale bar image and insert = 50μm. Values are shown as mean ± SEM.

Estrous cycles during lactation were significantly altered by deletion of Prlr in the forebrain, with all *Prlr*^lox/lox^/*Camk2a*^Cre^ mice showing a return to estrus between day 6 and day 10 of lactation (Figure 1E), while, as normal, estrus did not occur until day 20 (after weaning) in control *Prlr*^lox/lox^ mice (p = <0.0001, log-rank (Mantel-Cox) test, Figure 1E). No differences in litter weight gain from day 3 to day 8 of lactation were observed in either group (p = 0.3282, mixed analysis test, Supplementary Figure 4), indicating that the suckling stimulus that mice received and lactating itself was maintained in the absence of *Prlr* expression in *Camk2a* expressing neurons. Collectively, these data show that prolactin action in the brain is absolutely required for the lactation-induced suppression of kisspeptin expression and to maintain lactational infertility in mice.

In a separate cohort of *Prlr*^lox/lox^/*Camk2a*^Cre^ mice, pulsatile LH secretion was monitored in early lactation, prior to the return of estrous cycles. To rule out a potential role for progesterone in suppressing fertility during lactation (9,43,44), the progesterone receptor antagonist mifepristone (RU486), was administered to these mice in early lactation. Vehicle-treated *Prlr*^lox/lox^ control mice showed the expected near complete absence of pulsatile LH secretion during lactation (Figure 2A, B). In contrast, nearly all *Prlr*^lox/lox^/*Camk2a*^Cre^ mice showed a lack of the normal lactation-induced suppression of pulsatile LH secretion demonstrated by a significant increase in frequency of LH pulses compared to controls (effect of genotype p = 0.0024, two-way ANOVA with Tukey’s multiple comparisons test, Figure 2A, B). There was no effect of mifepristone on the pattern of LH secretion, suggesting that progesterone action is not required for the suppression of LH secretion (LH pulse frequency (interaction genotype x treatment p = 0.2807; treatment p = 0.8588; Figure 2B); mean LH levels (interaction genotype x treatment p = 0.8697; treatment p = 0.8586; Figure 2C); two-way ANOVA with Tukey’s multiple comparisons test). These data indicate that prolactin is the primary signal responsible for the suppression of pulsatile LH secretion during lactation in mice.

**Figure 2.**
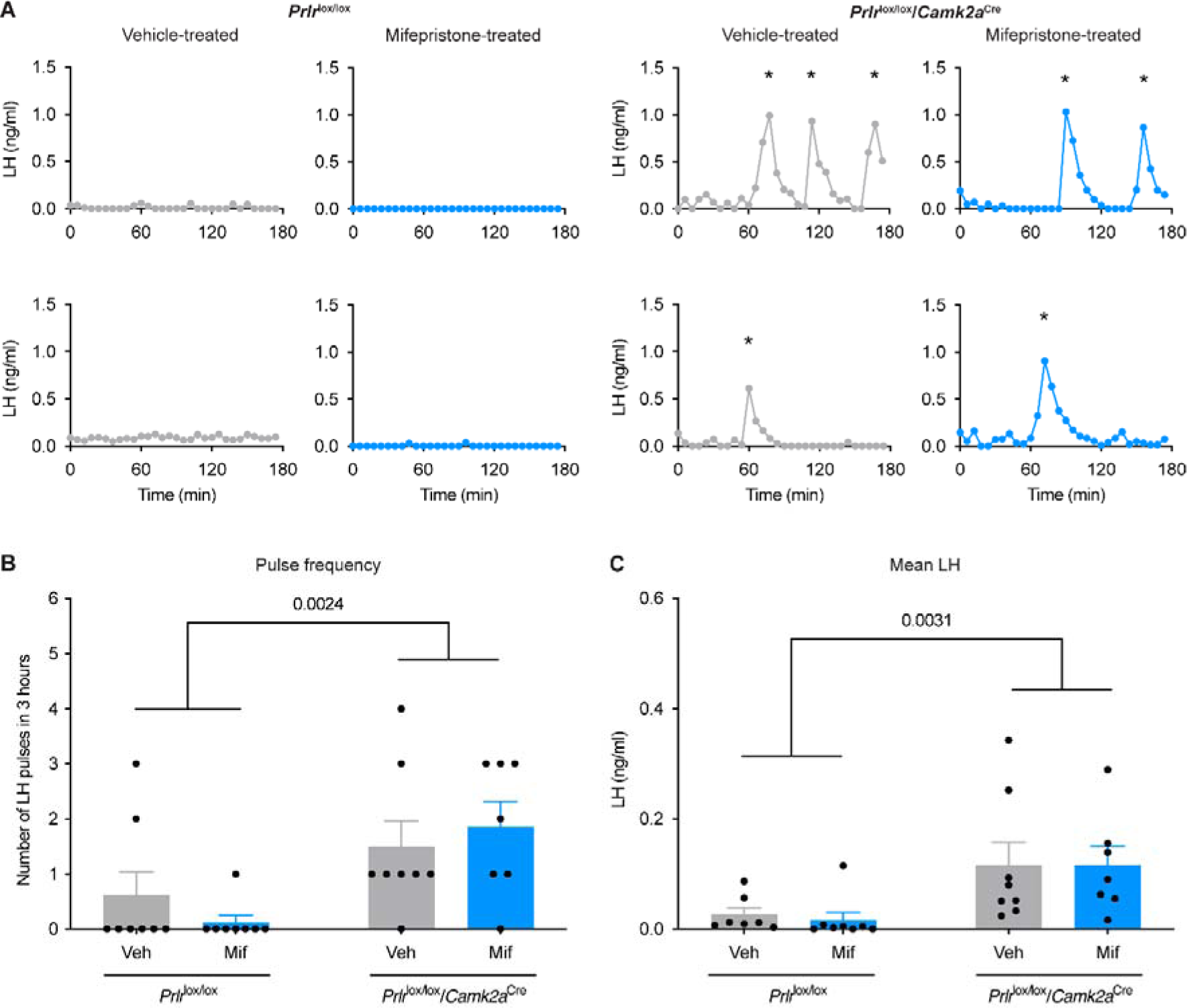
Prolactin action in the brain during lactation is necessary for the suppression of pulsatile LH secretion. Examples of pulsatile LH levels in the blood from lactating *Prlr*^lox/lox^ control and lactating *Prlr*^lox/lox^/*Camk2a*^Cre^ mice that have either been treated with vehicle (sesame oil, s.c., grey, veh) or 4mg/kg mifepristone (in sesame oil, s.c., blue, mif). Graphs show LH pulse frequency (B; interaction p = 0.2807, genotype p = 0.0024, state p = 0.8558), and mean LH levels (C; interaction p = 0.8697, genotype p = 0.0031, state p = 0.8586). Lactating vehicle-treated *Prlr*^lox/lox^ (n = 8), lactating mifepristone-treated *Prlr*^lox/lox^ (n = 8), lactating vehicle-treated *Prlr*^lox/lox^/*Camk2a*^Cre^ (n = 8), lactating mifepristone-treated *Prlr*^lox/lox^/*Camk2a*^Cre^ (n = 7). Asterisks indicate LH pulse peaks as detected by PULSAR Otago analysis. Two-way ANOVA followed by Tukey’s multiple comparisons test. Values are shown as mean ± SEM.

### Episodic activity of arcuate kisspeptin neurons is suppressed during pregnancy and most of lactation

To directly assess the role of prolactin in regulating kisspeptin neuron activity during lactation, GCaMP fibre photometry was undertaken to monitor real-time activity of arcuate kisspeptin neurons in freely behaving mice. We first undertook a longitudinal assessment of changes in kisspeptin neuronal activity by tracking individual *Kiss1*^Cre^ mice throughout pregnancy, lactation, and following weaning (Figure 3).

**Figure 3.**
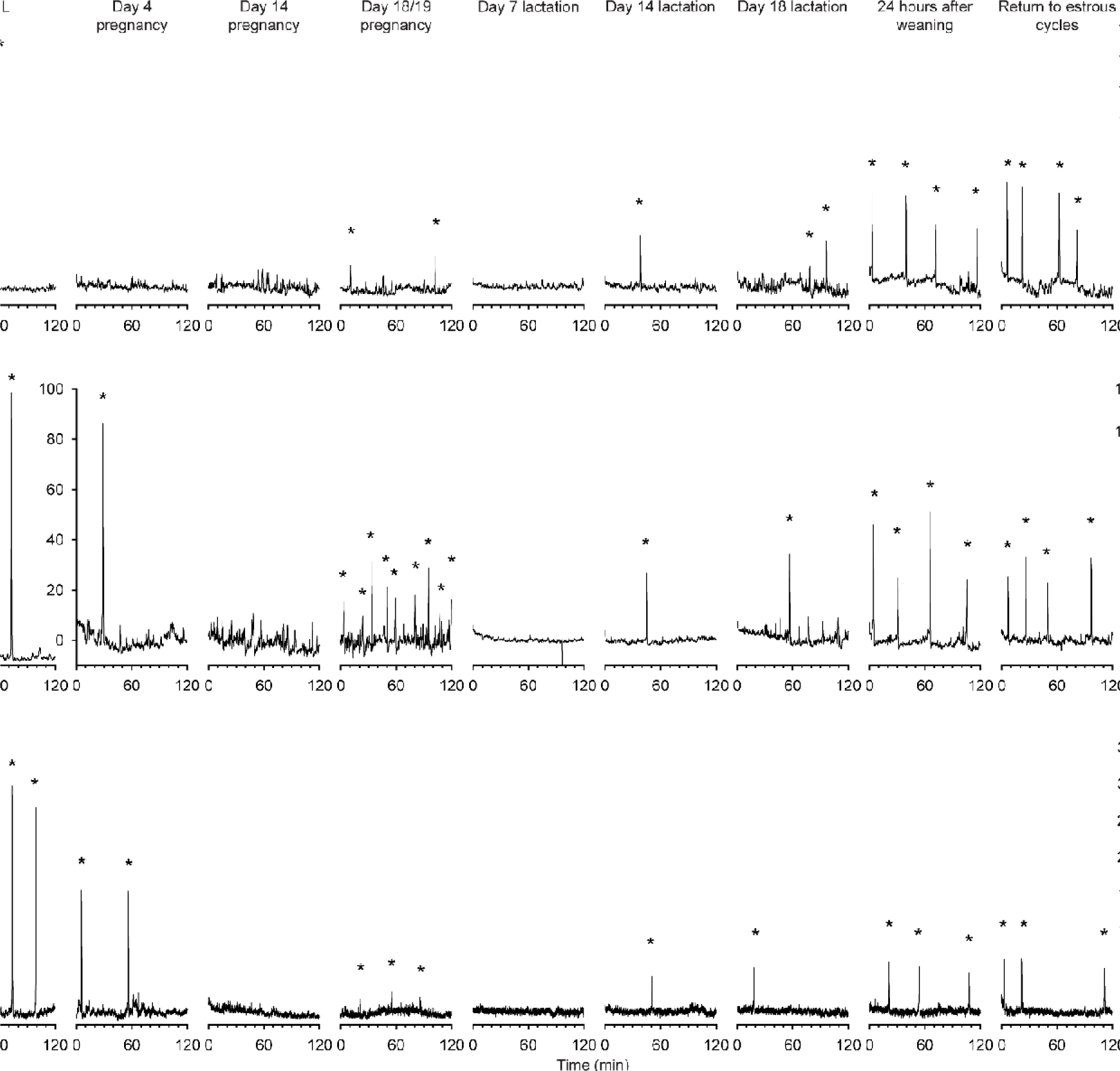
Longitudinal recordings of arcuate kisspeptin neuron GCaMP population activity throughout different reproductive states in the same *Kiss1*^Cre^ mice. Representative neuronal activity from three *Kiss1*^Cre^ mice throughout the NL (diestrus, nulligravid, non-lactating), pregnant, lactating, and post-weaning states. The time points monitored in order were: diestrus NL, day 4 pregnancy, day 14 pregnancy, day 18/19 pregnancy (overnight), day 7 lactation, day 14 lactation, day 18 lactation, 24 hours after weaning (day 22 postpartum), return to normal cycling following weaning (return to estrous cycles), and 10 days post ovariectomy (OVX). Asterisks indicate SEs. Note: change in y axes scale on all 3 OVX timepoints and mouse shown in dataset (B) from day 4 of pregnancy onwards.

Initially, GCaMP fibre photometry recordings were collected in the NL diestrous state, both with and without serial blood sampling to measure LH concentrations. As can be seen in Figure 4A, arcuate kisspeptin photometry recordings were characterized by discrete SEs of elevated intracellular calcium (indicative of synchronous activity of the kisspeptin population), with each SE consistently correlating to a single pulse of LH secretion in the minutes following (p = <0.0001, chi-squared test). These were observed to be at a similar rate to that described previously in diestrous mice using GCaMP photometry in a different *Kiss1* mouse line (9), with periodic SEs occurring about once per hour (1.250±0.250/hr; Figure 3 & 4B).

**Figure 4.**
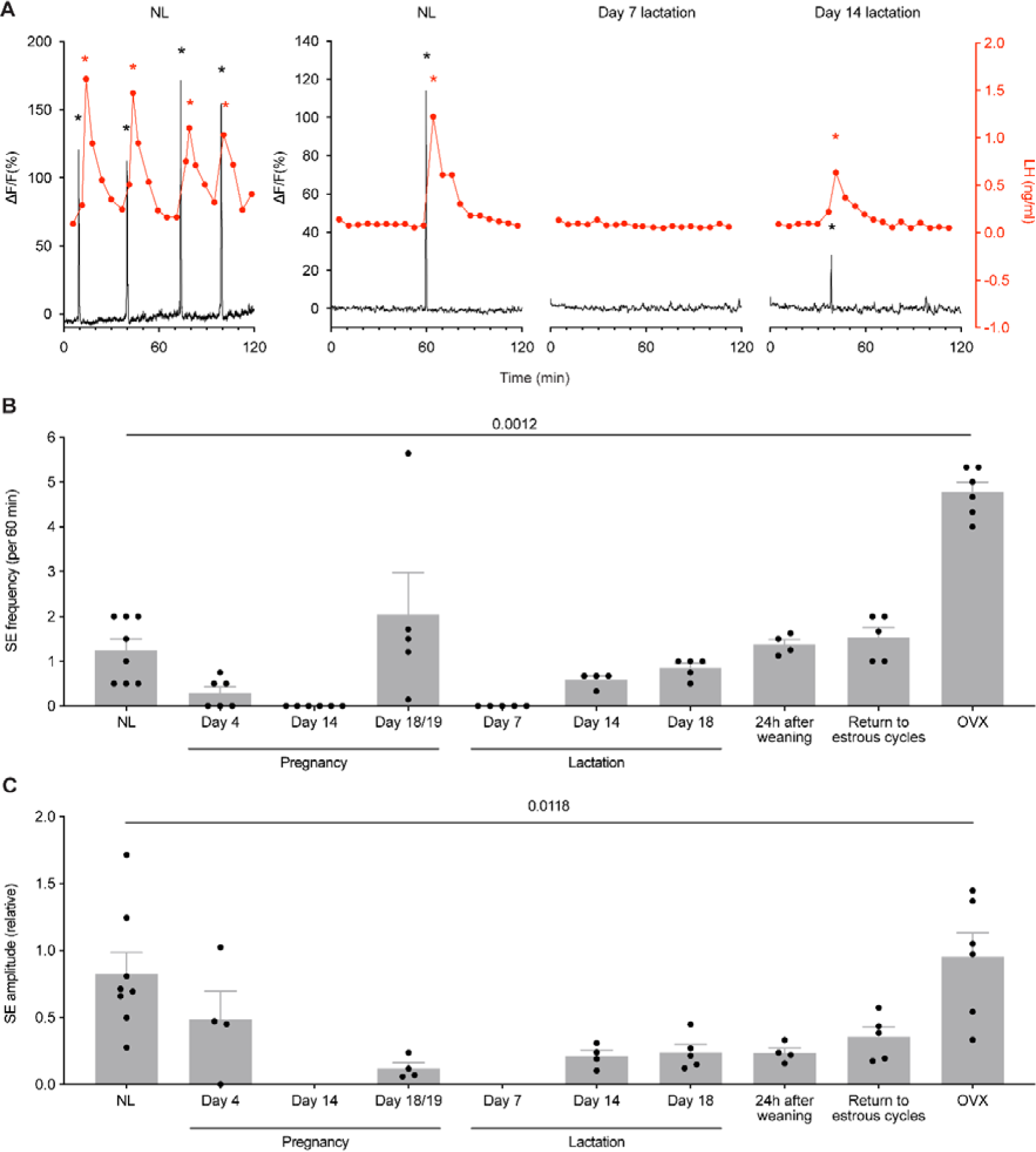
Synchronised Ca^2+^ events (SE) are consistently correlated with pulsatile LH secretion across different reproductive states in *Kiss1*^Cre^ mice. (A) When fibre photometry was paired with serial blood sampling for pulsatile LH secretion, the relationship between SEs and LH pulses was examined. Each time an SE was seen during a recording with blood sampling, pulsatile secretion of LH was also observed, with 100% correlation (p = <0.0001, chi-squared test; 73 out of 73 SEs observed led to an LH pulse). Representative examples of paired photometry and blood sampling are shown from the NL (diestrus, nulligravid, non-lactating) state, from day 7 lactation, and from day 14 lactation. (B) Quantitative analysis of SE frequency per hour across different reproductive states in *Kiss1*^Cre^ mice (p = 0.0012, mixed effect analysis (fixed type III) with Tukey’s multiple comparisons tests). (C) Quantitative analysis of relative SE amplitude of normalised ΔF/F across different reproductive states (p = 0.0118, mixed effect analysis (fixed type III) with Tukey’s multiple comparisons tests). Black asterisks indicate SEs, red asterisks indicate LH pulse peaks as detected by PULSAR Otago analysis. Values shown as mean ± SEM.

The activity of the arcuate kisspeptin population in *Kiss1*^Cre^ mice dynamically changed depending on the reproductive state of the mouse (p = 0.0012, mixed-effect analysis, Figure 3 & 4B). On day 4 of pregnancy, SE frequency had markedly decreased (0.297±0.136/hr; Figure 3 & 4B), indicating an early reduction in activity of arcuate kisspeptin neurons during pregnancy. By day 14 of pregnancy, no SEs were seen (0±0/hr; Figure 3 & 4B) and this was confirmed by a lack of pulsatile LH secretion (Supplementary Figure 5). In late pregnancy (day 18), neuronal activity was monitored for 14 hours, during which time low amplitude, SEs were unexpectedly observed at similar frequencies to NL levels (2.043±0.940/hr; Figure 3 & 4B-C). This unusual pattern of activity is illustrated in more detail in Figure 5 where, alongside the resurgence of low amplitude SEs, a marked increase in baseline activity was observed relative to recordings at other reproductive stages. This activity was reminiscent of the miniature SEs observed to be caused by activation of subgroups of arcuate kisspeptin cells using brain slice calcium imaging (11), but we are unable to resolve such SEs using the present methods. Since our aim was to continue longitudinal assessment of the arcuate kisspeptin population into lactation and weaning, mice were not disrupted by blood sampling immediately prior to parturition as we were concerned that this additional stressor might interfere with establishment of maternal behaviour. Hence, we are unable to report whether these low amplitude SEs and elevated baseline activity were associated with LH secretion.

**Figure 5.**
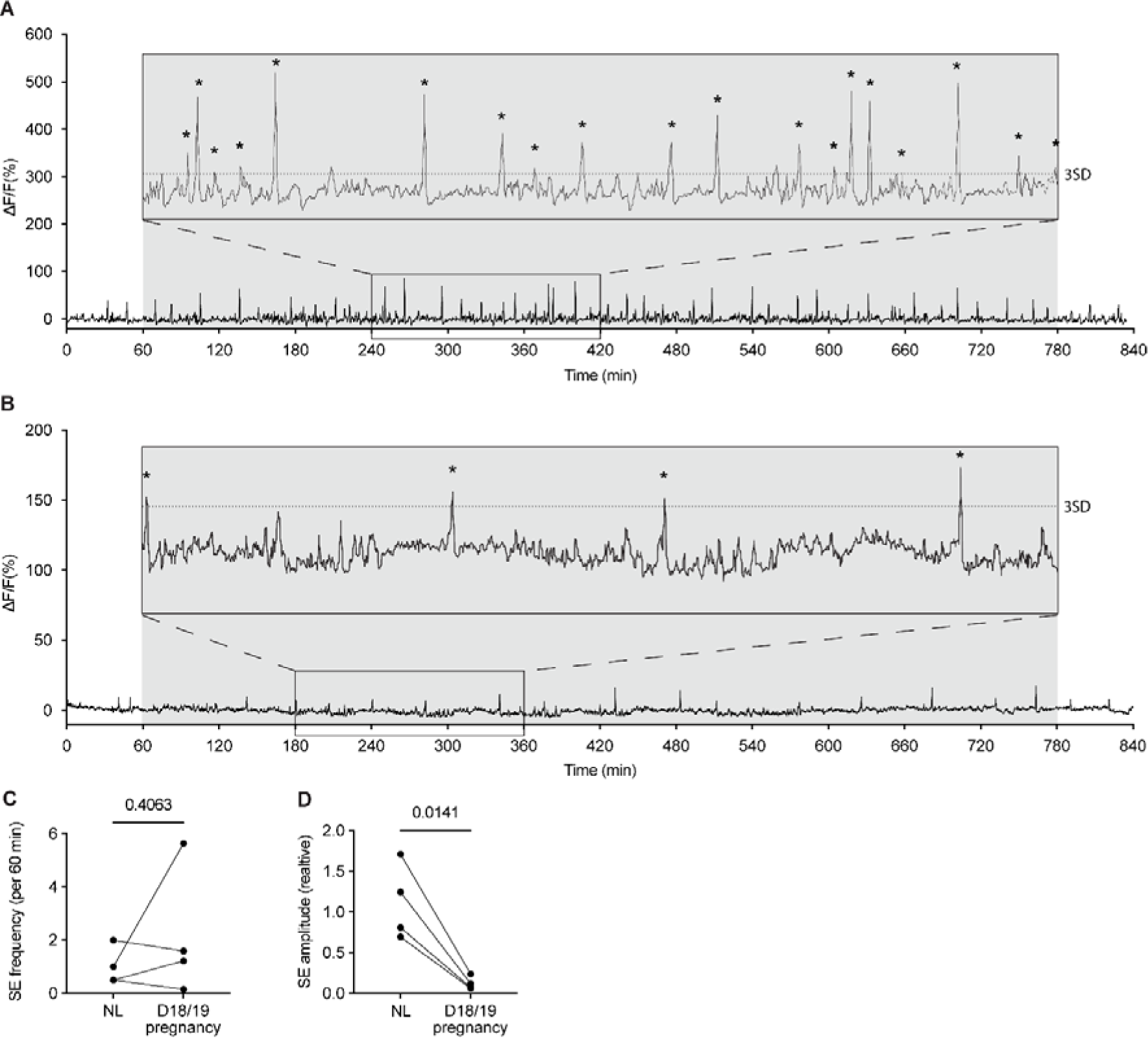
Activity of arcuate kisspeptin neurons on day 18 and 19 of pregnancy in *Kiss1*^Cre^ mice. Fibre photometry recordings of mice on the evening of day 18 of pregnancy (1800 hours) to the morning of day 19 of pregnancy (0800 hours) show low amplitude SEs. 3-hour section blown up for ease of viewing. (C) No difference was seen between frequency of SEs (per 60 minutes) in the NL (diestrus, nulligravid, non-lactating) versus day 18/19 pregnancy (p = 0.4063, paired two-tailed t test), however a significant decrease in relative SE amplitude is seen (D; p = 0.0141, paired two-tailed t test). Asterisks indicate SEs. Dotted line in insert of (A) and (B) indicates 3 standard deviations (3SD). Grey shaded region = lights off.

Evaluation of activity of arcuate kisspeptin neurons during lactation showed complete suppression of activity on day 7 of lactation with a corresponding absence of pulsatile LH secretion (0±0/hr; Figure 3 & 4B). This lactation-induced suppression of activity was partially relieved by day 14 of lactation (0.583±0.083/hr; Figure 3 & 4B), with low frequency SEs again corresponding to low frequency pulsatile LH secretion. Further increases in SE frequency were seen on day 18 of lactation (0.850±0.100/hr; Figure 3 & 4B), including an increase in baseline activity, similar to that seen in late pregnancy, and by 24 hours after weaning (day 22 postpartum) the frequency of SEs had returned to close to non-pregnant levels (1.375±0.114/hr; Figure 3 & 4B). SE frequency remained unchanged on the day of first diestrus following a return to estrous cycles after weaning (1.533±0.226/hr; Figure 3 & 4B). Ten days post-OVX, clusters of high amplitude arcuate kisspeptin population activity was observed (4.778±0.222/hr; Figure 3 & 4B), consistent with previous reports following OVX in nulliparous mice (45). Collectively, these observations show extensive, dynamic variation in the activity of the arcuate kisspeptin neuronal population associated with pregnancy and lactation.

### Mice with an arcuate kisspeptin neuron-specific deletion have premature reactivation of estrous cycles and neuronal activity in lactation

To determine whether the prolactin-induced suppression of estrous cycles and pulsatile LH secretion was specifically mediated by kisspeptin neurons, mice were generated with an arcuate-specific deletion of the Prlr from kisspeptin neurons (19). Similar to the data from *Prlr*^lox/lox^/*Camk2a*^Cre^ mice, there was early resumption of estrous cycles in *Prlr*^lox/lox^/*Kiss1*^Cre^ mice during lactation (63% showing estrus by day 10 of lactation and 83% by day 19 lactation) compared to *Prlr*^lox/lox^ controls (4% by day 19 lactation) (p = <0.0001, log-rank (Mantel-Cox) test, Figure 6A). No difference in litter weight gain during lactation was observed in either group indicating that suckling and/or lactation itself was not impaired (day 3-20 lactation weight gain, p = 0.6404, two-way ANOVA, Supplementary Figure 4H). *In vivo* GCaMP fibre photometry in *Prlr*^lox/lox^/*Kiss1*^Cre^ mice showed early reactivation of the arcuate kisspeptin population by day 5 of lactation (Figure 6B). This was accompanied by a clear return to estrus within this early lactation window in four out of five mice. These data demonstrate, in mice, that prolactin action specifically on arcuate kisspeptin neurons is responsible for maintaining suppression of those neurons, and thereby fertility, during lactation.

**Figure 6.**
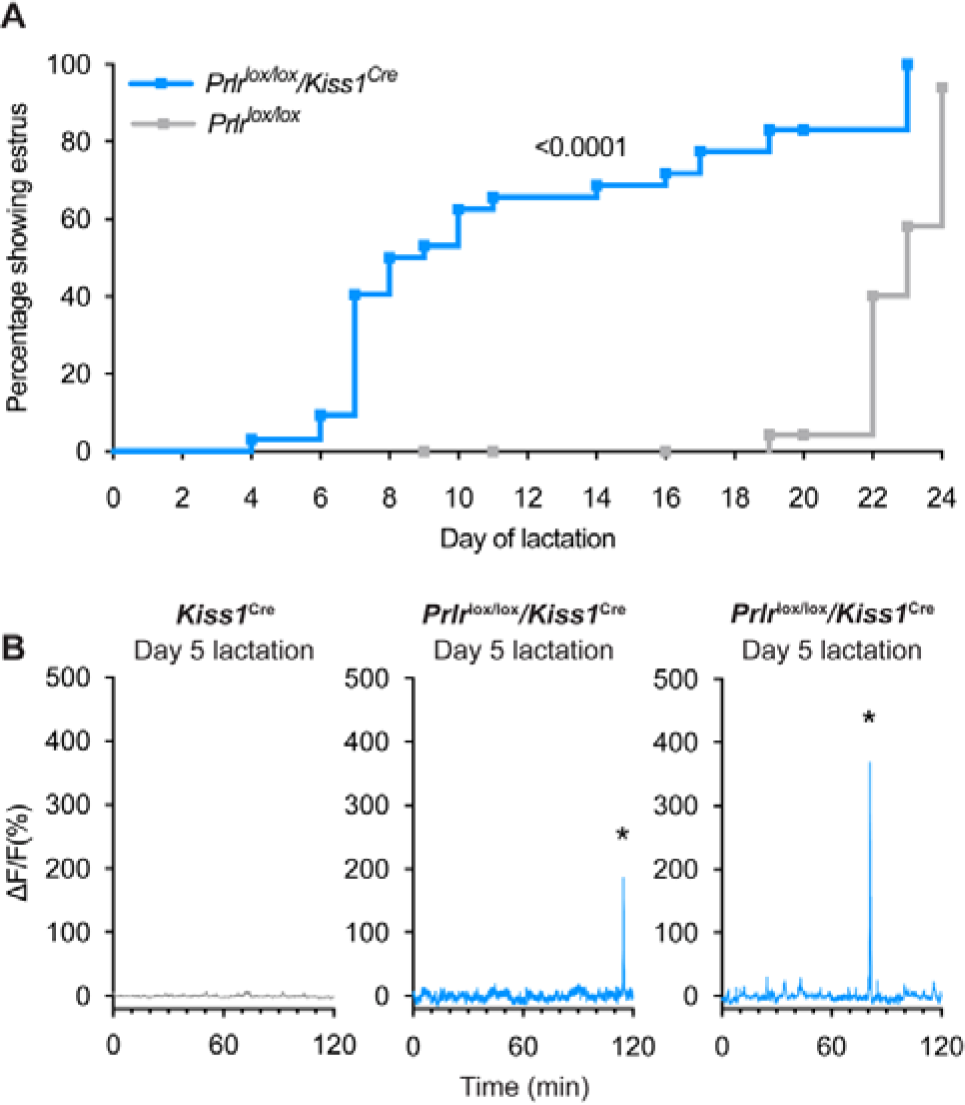
*Prlr*^lox/lox^/*Kiss1*^Cre^ mice do not undergo the normal period of lactational infertility and show early reactivation of arcuate kisspeptin neurons prior to estrus in lactation. (A) *Prlr*^lox/lox^/*Kiss1*^Cre^ mice resume estrous cycles significantly earlier (63% by day 10 of lactation, n = 32, blue) than *Prlr*^lox/lox^ controls (4% by day 19 of lactation, n = 30, grey) (p = <0.0001, Log-rank (Mantel-Cox) test). (B) Representative fibre photometry traces from day 5 of lactation from either a *Kiss1*^Cre^ control mouse or *Prlr*^lox/lox^/*Kiss1*^Cre^ mice. Mice with Prlr knocked out of arcuate kisspeptin neurons (*Prlr*^lox/lox^/*Kiss1*^Cre^) show SEs early in lactation. Asterisks indicate SEs.

## Discussion

We demonstrate here that prolactin action in arcuate kisspeptin neurons is necessary for the maintained suppression of fertility during lactation in mice. Neuron-specific Prlr deletion (*Prlr*^lox/lox^/*Camk2a*^Cre^) resulted in premature return to estrus in early lactation, even in the presence of ongoing suckling stimulus and the full metabolic consequences of milk production. Accompanying the resumption of estrus was an absence of the normal lactation-induced reduction in kisspeptin immunoreactivity (12,16,20). Pulsatile LH secretion was also observed on day 5 of lactation prior to the premature estrus when it would normally have been completely absent (46–48). To evaluate the specific role of kisspeptin neurons in mediating the prolactin-induced suppression of fertility, we have comprehensively mapped the activity of arcuate kisspeptin neurons throughout a full reproductive cycle: pregnancy, lactation, and after weaning in individual animals. The data show an immediate suppression of activity of arcuate kisspeptin neuronal activity during pregnancy, and this is maintained throughout most of lactation, apart from a brief window of reactivation immediately prior to parturition. Deleting Prlr specifically from arcuate kisspeptin neurons (*Prlr*^lox/lox^/*Kiss1*^Cre^) prevented the suppression of activity in early lactation, resulting in premature induction of episodic activation of kisspeptin neurons, and early onset of estrus. Combined, these data provide direct evidence that prolactin action on kisspeptin neurons is necessary for lactation-induced infertility in mice.

It is now well established that arcuate kisspeptin neurons form the GnRH “pulse generator”, and hence drive pulsatile release of GnRH from the hypothalamus and consequent pulsatile secretion of LH from the pituitary that is required for fertility (49–51). This is the first study to monitor activity of the arcuate kisspeptin neurons across different reproductive states in the same animal, and the data largely match previously described patterns of LH secretion (47,52). The frequency and dynamics of the SEs changed dramatically, initially due to the pregnancy-induced changes in ovarian hormones. The abrupt decrease in arcuate kisspeptin activity in early pregnancy is likely caused by rising levels of progesterone, known to profoundly suppress activity of arcuate kisspeptin neurons and LH secretion (9,43,44). Progesterone is elevated throughout pregnancy, gradually increasing until luteolysis and progesterone withdrawal occurs in the lead up to parturition (53–56). Interestingly, we observed a transient reactivation of the arcuate kisspeptin neurons in the night between days 18 and 19 of pregnancy. This was characterised by frequent, low amplitude SEs, and increased baseline activity that may represent the intermittent synchroniszed activity of small subsets of arcuate kisspeptin neurons that have not yet transitioned to full synchroniszation of the whole population (11). It seems likely that this pattern of activity is associated with progesterone withdrawal in late pregnancy and may be important in stimulating follicular growth leading up to a postpartum ovulation (57,58).

In early lactation, episodic activity of arcuate kisspeptin neurons was absent, with sporadic low-amplitude activity returning around day 14 of lactation. There was another period of increased baseline activity in late lactation, similar to that seen in late pregnancy, potentially representing a signature of reactivation of synchroniszed activity of the arcuate kisspeptin neurons. This increase in arcuate kisspeptin population activity during late lactation mirrors the increase in LH levels that has been reported as lactation progresses (59). Overall patterns of SE activity rapidly returned to normal diestrus levels soon after weaning (59). However, in the absence of Prlr in arcuate kisspeptin neurons, SE activity re-appeared as early as day 5 of lactation, even in the presence of ongoing suckling. These data clearly show that prolactin action in arcuate kisspeptin neurons is necessary to sustain lactational infertility in mice. The observed disruption of lactational infertility in *Prlr*^lox/lox^/*Kiss1*^Cre^ mice is particularly remarkable given that Prlr deletion is restricted to arcuate kisspeptin neurons in this model (19), and prolactin action on RP3V kisspeptin neurons (21,60) and on gonadotrophs in the pituitary gland (61–63) are unaffected.

The indispensable role for prolactin in mediating lactation-induced infertility in the mouse is surprising, given the consensus of much work in other species concluding that other factors may be more important (see (4,6,25,26,64)). This may reflect a level of redundancy amongst contributing factors across all species, including ovarian hormones, metabolic cues and neural inputs of suckling. Notably, the conditional deletion approach described here distinguishes prolactin action from the neurogenic effects of suckling without altering the process of lactation itself. Moreover, this approach avoids the potential confounding effects of using dopamine agonists to suppress prolactin (28,29), given that dopamine can directly inhibit GnRH neuronal activity (65).

While the effects of widespread neuronal deletion (*Prlr*^lox/lox^/*Camk2a*^Cre^) on fertility were largely recapitulated by the arcuate kisspeptin-specific model (*Prlr*^lox/lox^/*Kiss1*^Cre^), it was apparent that the global deletion was more effective at inducing the return to estrus during lactation (in 100% of *Prlr*^lox/lox^/*Camk2a*^Cre^ mice by day 10), compared to the arcuate kisspeptin-specific model (63% by day 10, and 83% by day 19). This may be due to the absence of lactation-induced suppression of *Kiss1* expression in RP3V kisspeptin neurons of *Prlr*^lox/lox^/*Camk2a*^Cre^ mice. Similarly, we cannot rule out the possibility that other populations of prolactin-sensitive neurons, such as GABA or dopamine neurons (60), may contribute to suppressing estrous cycles during lactation.

Nevertheless, our data collectively provide strong evidence that prolactin action on arcuate kisspeptin neurons is the primary factor mediating lactation-induced infertility in mice. Given that hyperprolactinemia induces infertility in many other mammalian species, including humans (21,22,66–75), it is possible that a conserved mechanism could be contributing to lactational infertility in these species.

## Author Contributions

ECRH: Designed research, completed research, analysed data, wrote the first draft of the manuscript, edited the manuscript.

SRL: Designed research, completed research, analysed data, edited the manuscript.

JC: Completed research, analysed data, edited the manuscript.

HJM: Completed research, edited the manuscript.

UB: Provided key resources, edited the manuscript.

AEH: Designed research, analysed data, edited the manuscript.

RSEB: Designed research, analysed data, edited the manuscript.

DRG: Designed research, provided key resources, analysed data, edited the manuscript.

## Acknowledgments

We would like to acknowledge the research assistance of Zin Khant-Aung, genotyping by Pene Knowles, and Dr Joon Kim for assistance with analysis of fibre photometry data.

Financial support: This work was supported by Health Research Council of New Zealand (grant number: 21-560) and the Lions Club of Dunedin South - administered by Perpetual Guardian (Otago Medical Research Foundation).

## Supplementary data

**Supplementary Figure 1.**
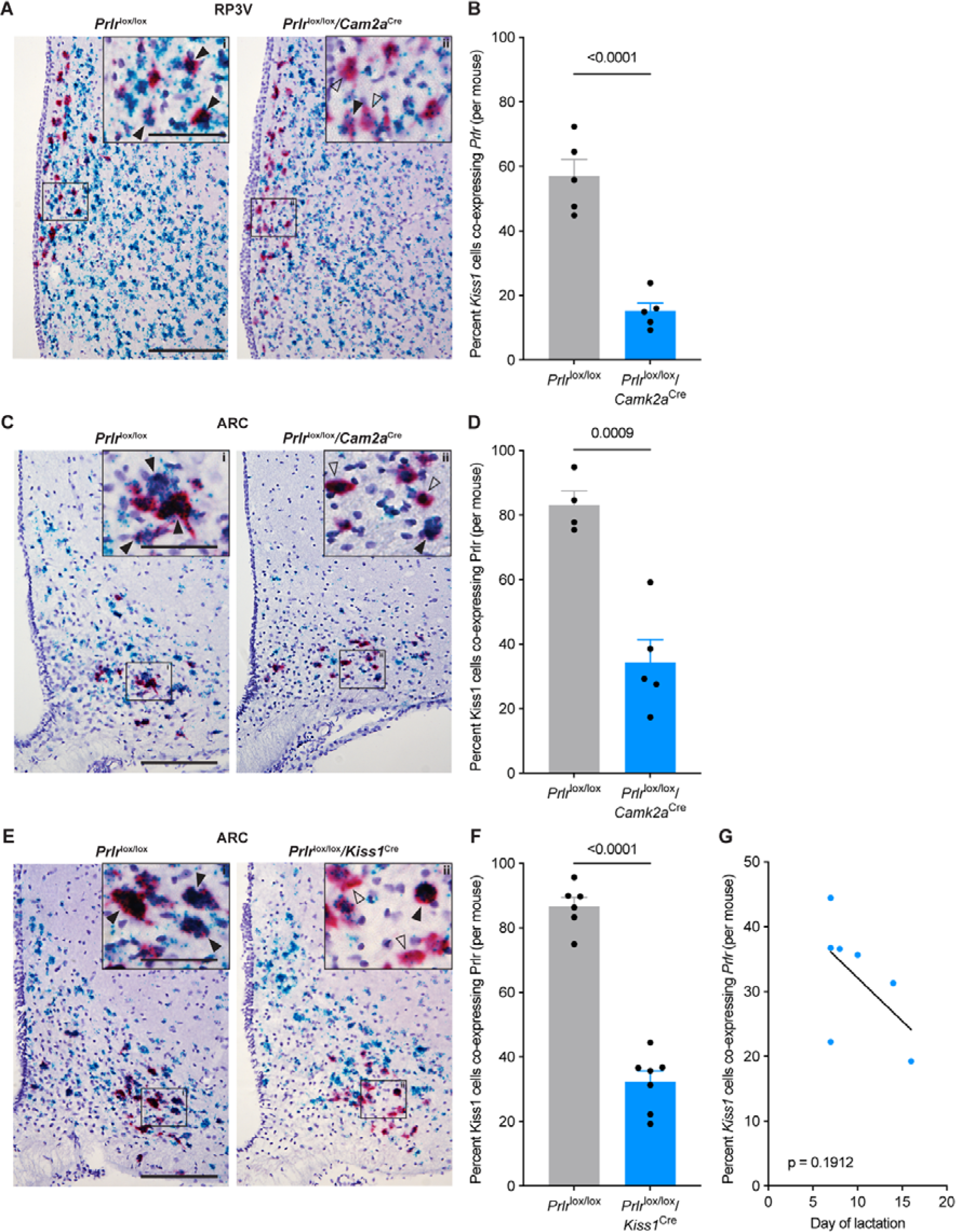
Proportion of kisspeptin neurons showing *Prlr* deletion using RNAscope in either *Prlr*^lox/lox^/*Camk2a*^Cre^ or *Prlr*^lox/lox^/*Kiss1*^Cre^ mice. Representative photomicrographs showing RNAscope labelling for *Prlr* (blue) and *Kiss1* (red) in the rostral periventricular region of the third ventricle (RP3V, A) or arcuate nucleus (ARC, C, E), in either intact (*Prlr*^lox/lox^ control n = 5, *Prlr*^lox/lox^/*Camk2a*^Cre^, n = 5, A) or ovariectomised (OVX; *Prlr*^lox/lox^ control n = 4, *Prlr*^lox/lox^/*Camk2a*^Cre^ n = 5, C; *Prlr*^lox/lox^ control n = 6, *Prlr*^lox/lox^/*Kiss1*^Cre^ n = 7, E) mice. Compared to *Prlr*^lox/lox^ control mice, *Prlr*^lox/lox^/*Camk2a*^Cre^ mice show a significant decrease in the percentage of *Kiss1*-expressing cells co-expressing *Prlr* in both the RP3V (B; p = <0.0001) and arcuate nucleus (D; p = 0.0009) (unpaired two-tailed t tests). (F) A significant decrease in the percentage of *Kiss1*-expressing cells co-expressing *Prlr* was seen in *Prlr*^lox/lox^/*Kiss1*^Cre^ compared to *Prlr^l^*^ox/lox^ controls (p = <0.0001, unpaired two-tailed t test). (G) No correlation was found between percentage of *Kiss1* cells co-expressing with *Prlr* and the day of estrus return during lactation (p = 0.1912, simple linear regression). Solid black arrows = doubled labelled cells expressing both *Kiss1* and *Prlr*; black outlined arrows = *Kiss1* cells with sparse or no co-labelling for *Prlr*. Scale bar = 150μm, insert = 60μm. Values are shown as mean ± SEM.

**Supplementary Figure 2.**
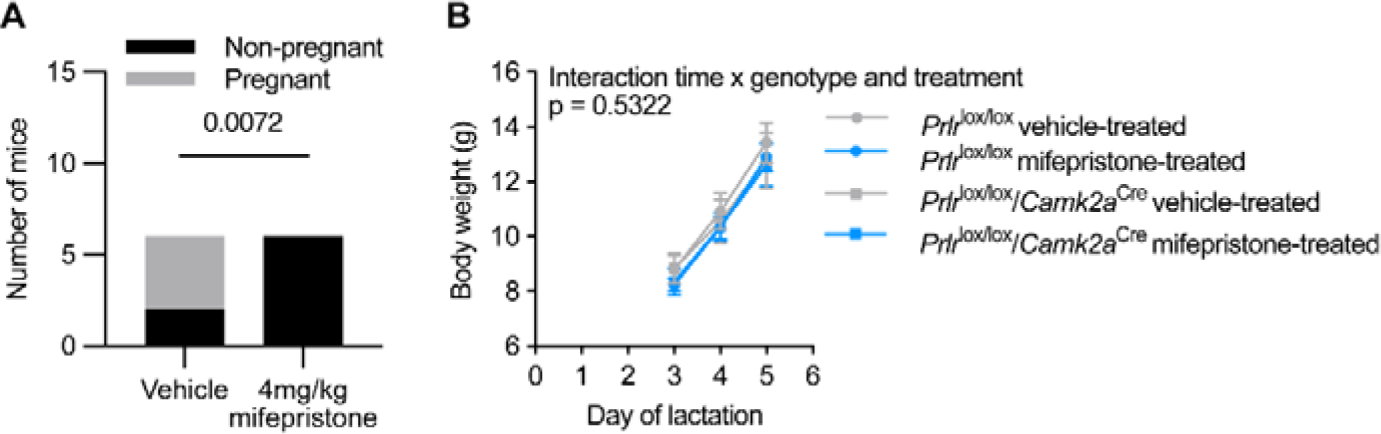
Mifepristone dose response and effect on litter weight gain. (A) Mifepristone dose response trial showing dose of 4mg/kg was sufficient to terminate pregnancy in all mice (p = 0.072, chi-squared test, n = 6 both groups). (B) Mifepristone or vehicle treatment had no effect on litter weight gain from day 3 to day 5 of lactation (interaction of time x genotype & treatment p = 0.5322; time p = <0.0001; genotype and treatment p = 0.8811; subject p = <0.0001; two-way ANOVA). Values are shown as mean ± SEM.

**Supplementary Figure 3:**
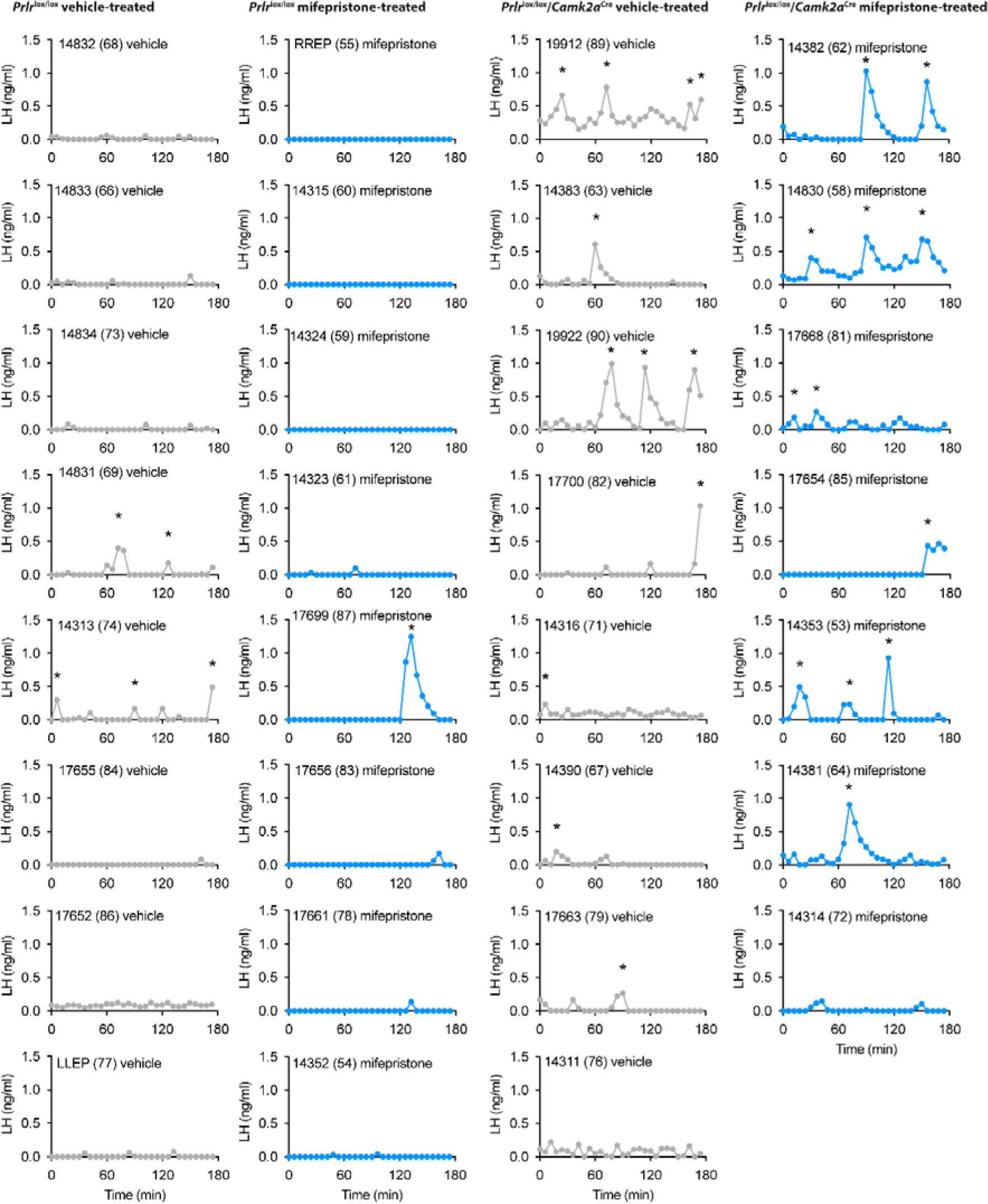
Pulsatile LH secretion profiles of *Prlr*^lox/lox^/*Camk2a*^Cre^ mice and *Prlr*^lox/lox^ controls following vehicle or mifepristone treatment. Individual LH pulse data from lactating *Prlr*^lox/lox^ control and *Prlr*^lox/lox^/*Camk2a*^Cre^ mice treated with either vehicle (sesame oil, s.c., grey) or 4mg/kg mifepristone (in sesame oil, s.c., blue). Lactating vehicle-treated *Prlr*^lox/lox^ (n = 8), lactating mifepristone-treated *Prlr*^lox/lox^ (n = 8), lactating vehicle-treated *Prlr*^lox/lox^/*Camk2a*^Cre^ (n = 8), lactating mifepristone-treated *Prlr*^lox/lox^/*Camk2a*^Cre^ (n = 7). Asterisks indicate LH pulse peaks as detected by PULSAR Otago analysis.

**Supplementary Figure 4.**
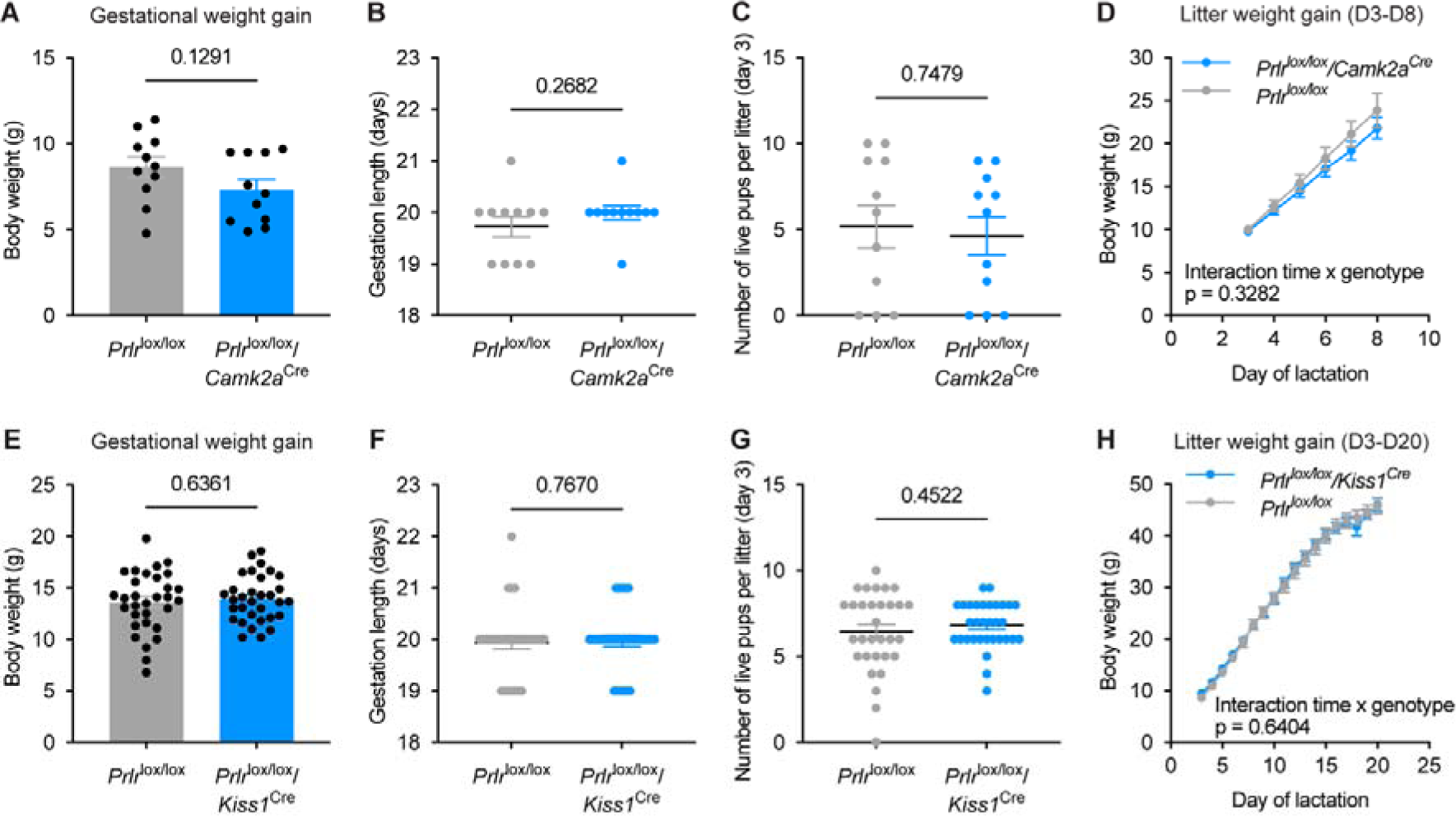
Gestational and maternal phenotyping of *Prlr*^lox/lox^/*Camk2a*^Cre^ and *Prlr*^lox/lox^/*Kiss1*^Cre^ mice and their respective *Prlr*^lox/lox^ controls. (A-H) Data show; gestational weight gain (A; *Prlr*^lox/lox^ n = 11, *Prlr*^lox/lox^/*Camk2a*^Cre^ n = 11; E; *Prlr*^lox/lox^ n = 31, *Prlr*^lox/lox^/*Kiss1*^Cre^ n =27), gestation length (B; *Prlr*^lox/lox^ n = 11, *Prlr*^lox/lox^/*Camk2a*^Cre^ n = 11; F; *Prlr*^lox/lox^ n = 31, *Prlr*^lox/lox^/*Kiss1*^Cre^ n =27), number of live pups on day 3 of lactation (C; *Prlr*^lox/lox^ n = 11; *Prlr*^lox/lox^/*Camk2a*^Cre^ n = 11; G; *Prlr*^lox/lox^ n = 31, *Prlr*^lox/lox^/*Kiss1*^Cre^ n =27), and litter weight gain between day 3-8 of lactation (D; *Prlr*^lox/lox^ n = 8; *Prlr*^lox/lox^/*Camk2a*^Cre^ n = 8) or day 3-20 of lactation (H; *Prlr*^lox/lox^ n = 22, *Prlr*^lox/lox^/*Kiss1*^Cre^ n = 20). There were no differences in any of these parameters (A, C, E, G, unpaired two-tailed t test; B, F, Mann Whitney test; D, H, repeated measures mixed effect analysis, fixed effects (type III) with Šídák’s multiple comparisons test. Grey = *Prlr*^lox/lox^; blue in top row A-D = *Prlr*^lox/lox^/*Camk2a*^Cre^, blue in bottom row E-H = *Prlr*^lox/lox^/*Kiss1*^Cre^. N numbers of *Prlr*^lox/lox^/*Kiss1*^Cre^ mice and respective *Prlr*^lox/lox^ controls changed due to: *Prlr*^lox/lox^ (n = 5), and *Prlr*^lox/lox^/*Kiss1*^Cre^ (n = 5) mice euthanised prior to day 20 lactation due to COVID-19 lockdown; *Prlr*^lox/lox^/*Kiss1*^Cre^ (n = 2) showed estrus and therefore underwent blood sampling (with respective *Prlr*^lox/lox^ controls (n =2)). All these mice were euthanised 2 hours following blood sampling; a *Prlr*^lox/lox^ dams’ (n =1) litter losing weight and being excluded from subsequent analysis. Values are shown as mean ± SEM.

**Supplementary Figure 5.**
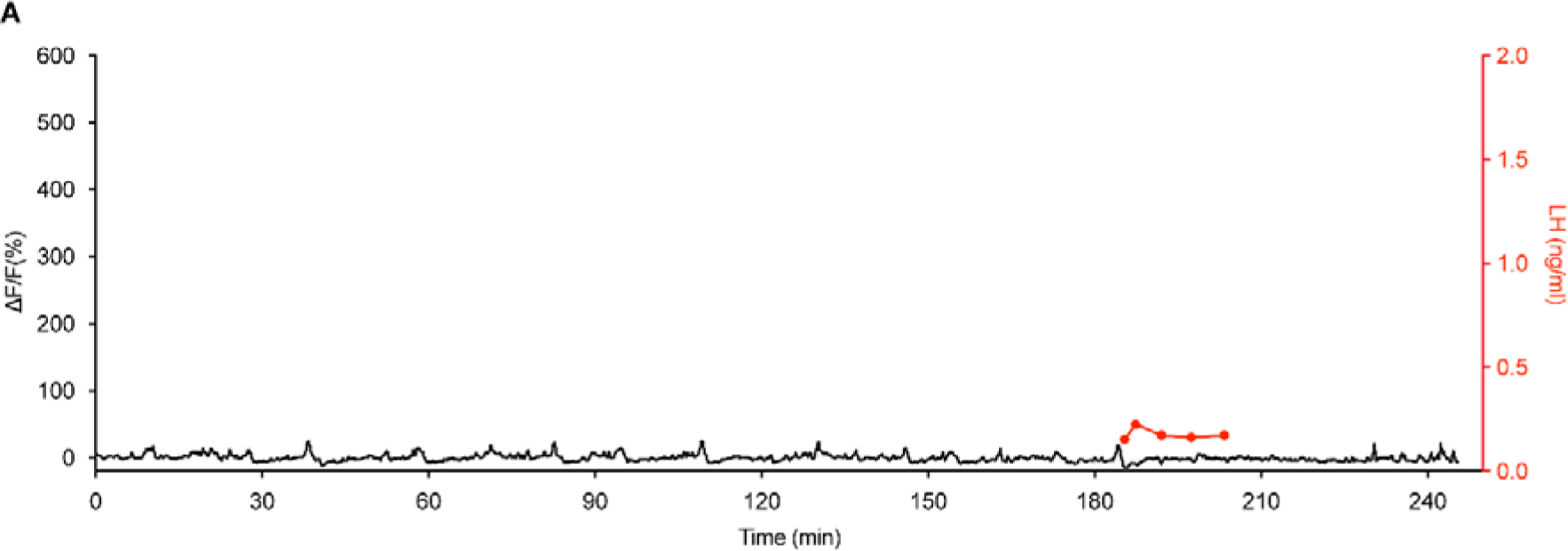
Miniature synchronised event (SE)-like activity on day 14 of pregnancy in *Kiss1*^Cre^ mice does not result in pulsatile LH secretion. (A) Paired fibre photometry and blood sampling from representative mouse on day 14 of pregnancy showing miniature SE-like activity have no significant effect on pulsatile LH secretion (red).

**Supplementary Table 1.**
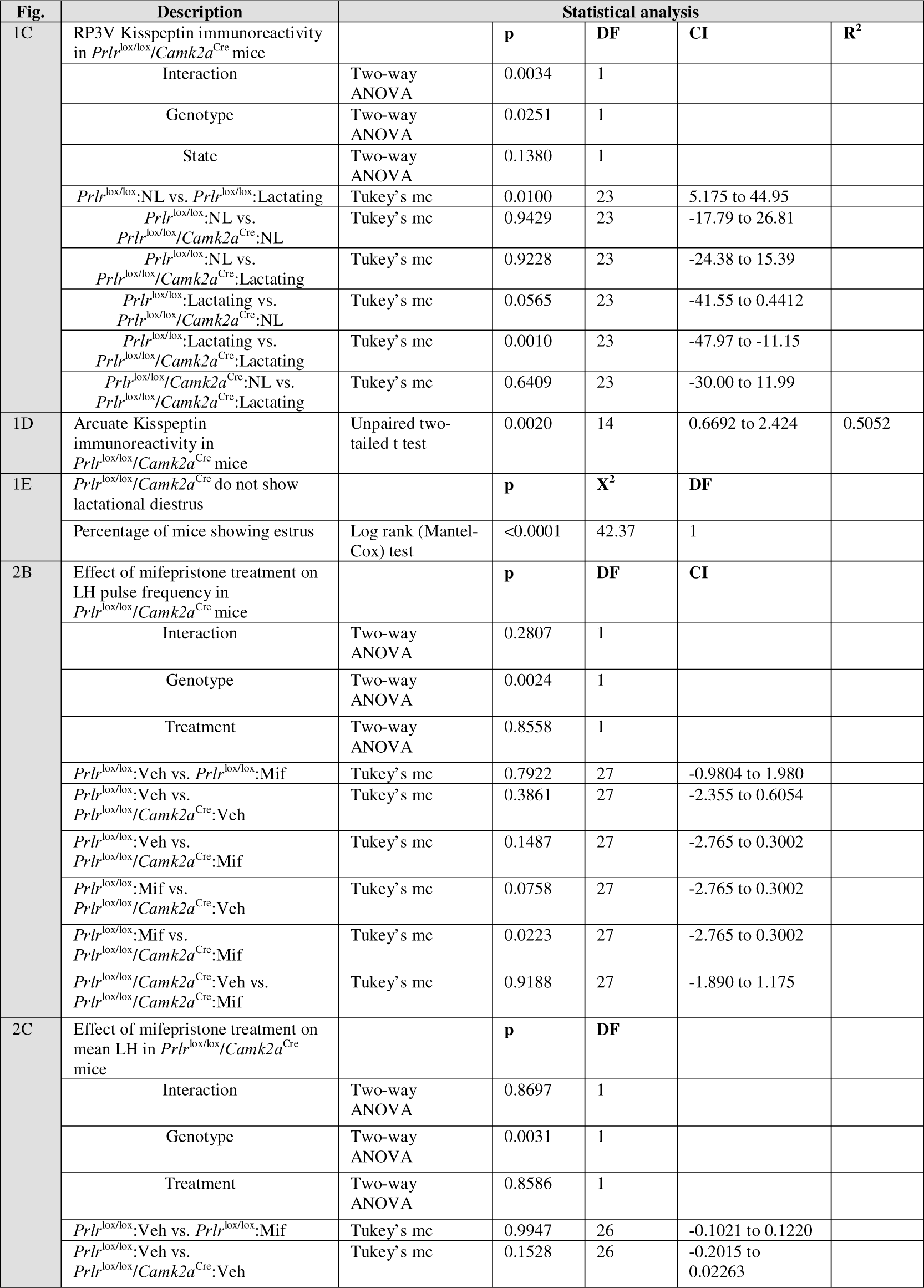
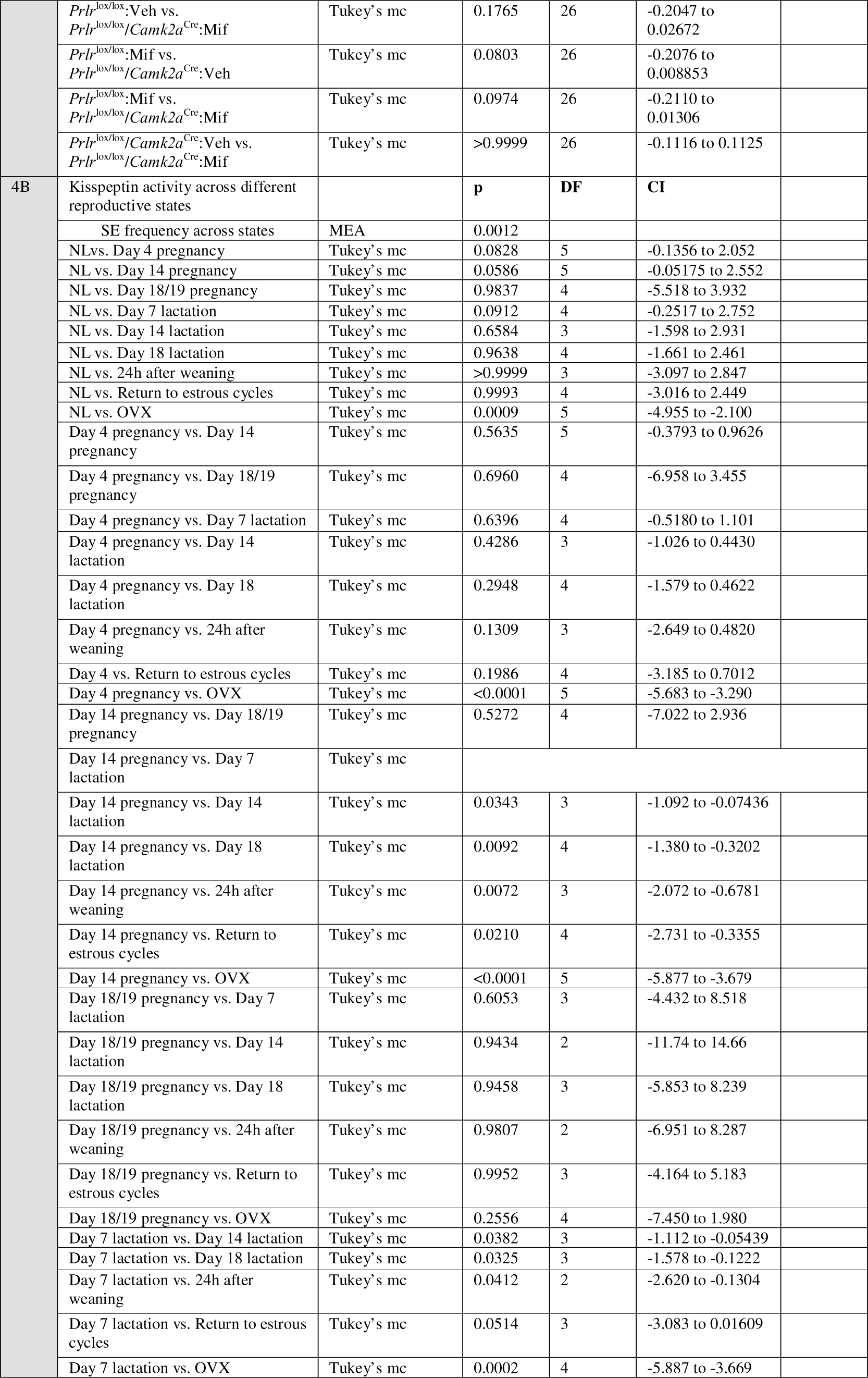
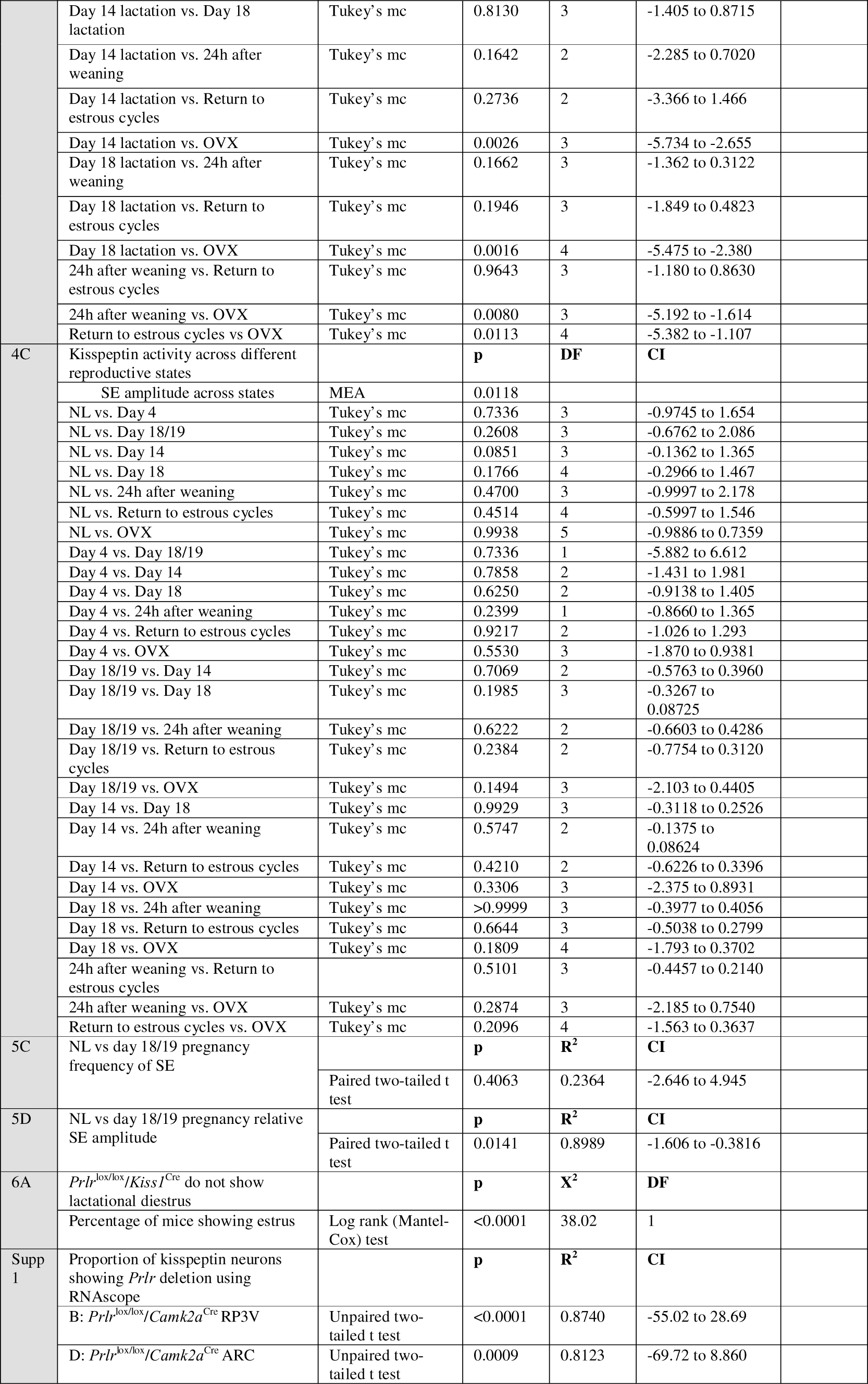
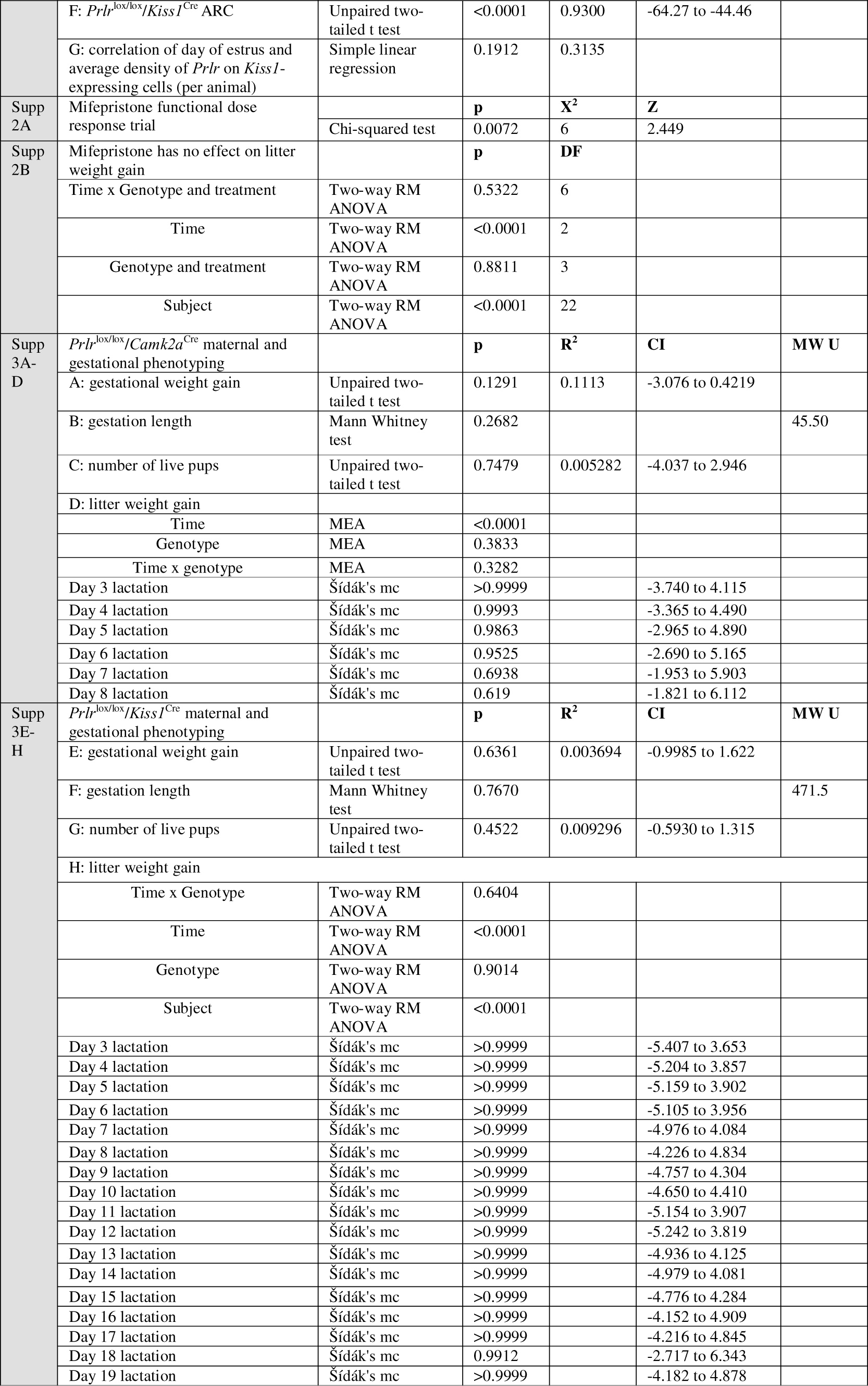
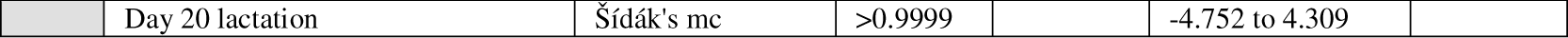
Statistics table. Abbreviations for tables below: DF = degrees of freedom; mc = multiple comparison; CI = 95% confidence interval; MW U = Mann Whitney U; MEA = Mixed effect analysis (fixed effects (type III)), RM = repeated measures. Supp = supplementary data figure.

